# Hippocampal ripples initiate cortical dimensionality expansion for memory retrieval

**DOI:** 10.1101/2025.04.22.649929

**Authors:** Casper Kerrén, Sebastian Michelmann, Christian F Doeller

## Abstract

How are past experiences reconstructed from memory? Learning is thought to compress external inputs into low-dimensional hippocampal representations, later expanded into high-dimensional cortical activity during recall. Hippocampal ripples, brief high-frequency bursts linked to retrieval, may initiate this expansion. Analysing intracranial EEG data from patients with pharmaco-resistant epilepsy during an episodic memory task, we found that cortical dimensionality increased following ripple events during correct, but not incorrect, retrieval. This expansion correlated with faster reaction times and reinstatement of the target association. Crucially, hippocampal theta and cortical gamma phase-amplitude coupling emerged after ripples but before cortical expansion, suggesting a mechanism for ripple-driven communication. Ripple events also marked the separation of task-relevant variables in cortical state space, revealing how hippocampal output reshapes the geometry of memory representations to support successful recall.

## Introduction

Episodic memory allows us to store detailed records of past experiences and consciously reconstruct those experiences at later points in time ^1^. Like any computational system though, the human brain operates with finite resources ^2^. To cope with these constraints, efficient encoding and retrieval of memories is thought to rely on compression and expansion of neural representations ^3–6^. During encoding, environmental information flows through cortical and subcortical regions to the hippocampus, where memories are initially stored ^7,8^. During retrieval, internal or external cues trigger the hippocampus to detect matches with stored traces ^9^. A partial match initiates pattern completion, leading to memory reactivation and reconstruction in cortical networks ^10,11^. Yet, how hippocampal pattern completion gives rise to cortical reinstatement remains poorly understood.

Recent studies have begun to conceptualise this process as shifts in the geometric relationships of points in neural state space, a framework that has found broader applications in studies of decision-making and working memory ^12–19^. We hypothesise that memory retrieval might likewise involve a transformation in dimensionality, such that low-dimensional hippocampal representations are expanded into a higher-dimensional cortical state, allowing mnemonic information to be decoded for successful recall ^5,10,20–22^. Preliminary evidence supports this hypothesis, showing a shift from semantic to perceptual representations along the ventral visual stream during retrieval ^23–27^. However, whether and how the hippocampus might initiate this dimensionality expansion is unclear.

One potential means to drive cortical representational expansion is through hippocampal ripples. Ripples are known to coordinate the transfer of compressed representations and changes of brain-wide functional connectivity during offline periods in rodents ^28–32^. In humans, an increase of hippocampal ripple rates precedes episodic memory recall ^33–39^ and neocortical reinstatement of previously encoded memories consistently follows hippocampal ripple events ^33–36^. In parallel, theoretical accounts and empirical evidence have highlighted cross-frequency interactions, particularly theta-gamma-phase-amplitude coupling (TG-PAC), as a mechanism for coordinating long-range communication ^40–46^. While TG-PAC has been widely studied during mnemonic processing particularly within hippocampus ^47–55^, its role during retrieval, and specifically in mediating ripple-initiated cortical transformations, remains largely unexplored.

We hypothesise that the reinstatement of information in cortical regions is supported by a ripple-based mechanism, where compressed representations are expanded via PAC-based connectivity between hippocampus and cortex. To test these hypotheses, we analysed intracranial data from 12 patients with pharmaco-resistant epilepsy as they performed an associative recognition memory task (Figs. 1a and b, Supplementary Figs. 1-2). We asked whether ripple events were related to both an increase in cortical representational dimensionality and TG-PAC between hippocampus and cortex, linking local hippocampal dynamics to global cortical transformations during successful episodic retrieval (Fig. 1c).

**Fig. 1.**
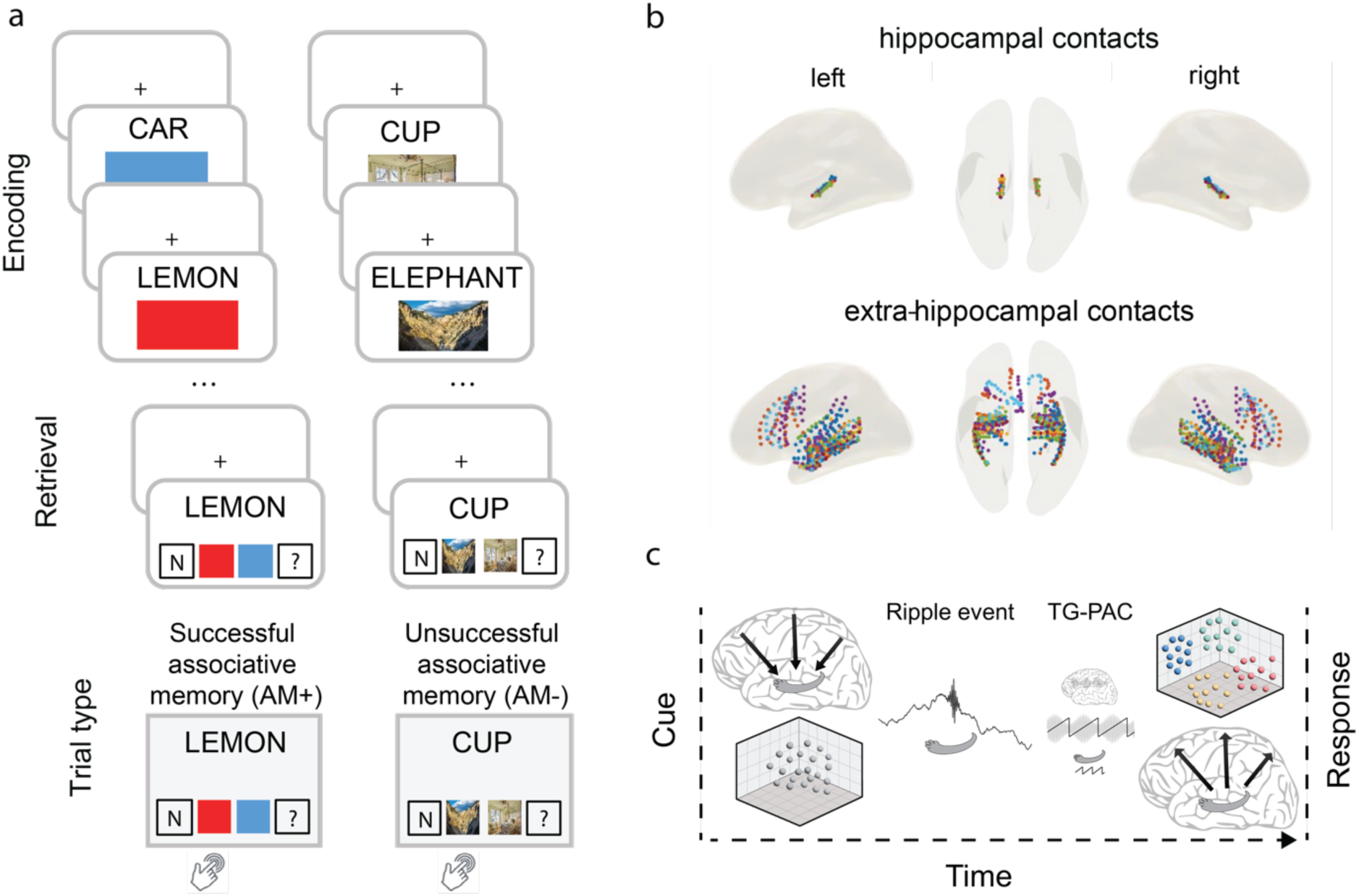
Paradigm, implantation scheme, hypothesis. **a** During encoding, participants associated a noun either with one of two colours (*left*) or one of two scenes (*right*; alternating across runs), and indicated whether the combination was plausible or implausible. During retrieval, participants were prompted with the noun alongside response options, yielding either successful recognition and associative memory (AM+) or successful recognition, but unsuccessful associative memory (AM-; combining incorrect and ‘don’t know’ [‘?’] responses). ‘N’ stands for new items. **b** Hippocampal (*top*) and extra-hippocampal (*bottom*) iEEG contacts included in the analyses, highlighted in different colours for each participant. **c** Schematic depiction of the hypothesis: During encoding, the representation of an event is stored in hippocampus as a lower-dimensional pointer to the higher-dimensional representation in cortical areas. During retrieval, hippocampal ripples initiate a process that initiates a hippocampal-cortical connectivity through theta-gamma phase-amplitude coupling (TG-PAC), leading to the low-dimensional representation being expanded in cortical areas, which is linked to behavioural memory performance.

### Behavioural results

Participants were on average good at recognising old items (75% hits ± 3.67%, mean ± SE), as well as correctly rejecting new items (79.83% ± 4.87%). Within hits, the experiment yielded a balanced amount of successful and unsuccessful associative memory trials (AM+: 49.95% ± 5.12%; AM-: 50.05% ± 5.12%; *t*(1,11) = -.01, *p* = .992). AM- trials contained trials both when participants indicated that the association was old, but they chose the wrong association, and when they indicated that the association was old, but they couldn’t remember [‘?’]. Associative incorrect proportion of hits was 16.28% (± 2.40%), whereas don’t know responses (‘?’) was 33.77% (± 7.07%). On trials where they either picked the correct or incorrect association (AM+ and incorrect), associative accuracy was markedly above the 50% guessing level (76% ± 2%, range = 64% - 86%; paired-samples t-test against chance: *t*(11) = 12.76, *p* > .01), demonstrating that participants did not simply guess between the two options when they committed to an associative decision. Response latencies for AM+ trials were faster (1.90 sec ±.12 sec) than for AM- trials (2.06 sec ± .12 sec; *t*(11) = −2.68, *p* = .021).

### Greater ripple-density for successful retrieval

From 72 hippocampal contacts (6.0 ± .64 mean ± SE, range 2-10) across 12 participants, on average, 787.33 hippocampal ripples (±145.97, range 220-2048) per participant were detected during all retrieval trials, after excluding false positives, with a spectral mean peak at 89.17 Hz (± .57 Hz) (Fig. 2a; Supplementary Fig. 1 for participant-specific ripple plots, and Supplementary Table 1 for breakdown per participant). Previous work has shown that hippocampal ripples are strongly phase-locked to low-frequency activity (e.g., ^56,57^). Therefore, we next examined the relationship between ripple timing and delta-band (.5-2 Hz) phase. Across all detected ripples, we observed a clear non-uniform clustering of phases where ripples preferentially occurred at a mean delta phase of 37° (Rayleigh test: *z* = 12.85, *p* < .01, Rayleigh statistic against a surrogate distribution generated by shuffling ripple times within trials, *p* =.013; inset Fig. 2a), consistent with previous human intracranial findings. These results indicate that ripples in our dataset tend to occur on the descending phase of slow hippocampal oscillations, suggesting that ongoing delta activity helps gate ripple initiation.

**Fig. 2.**
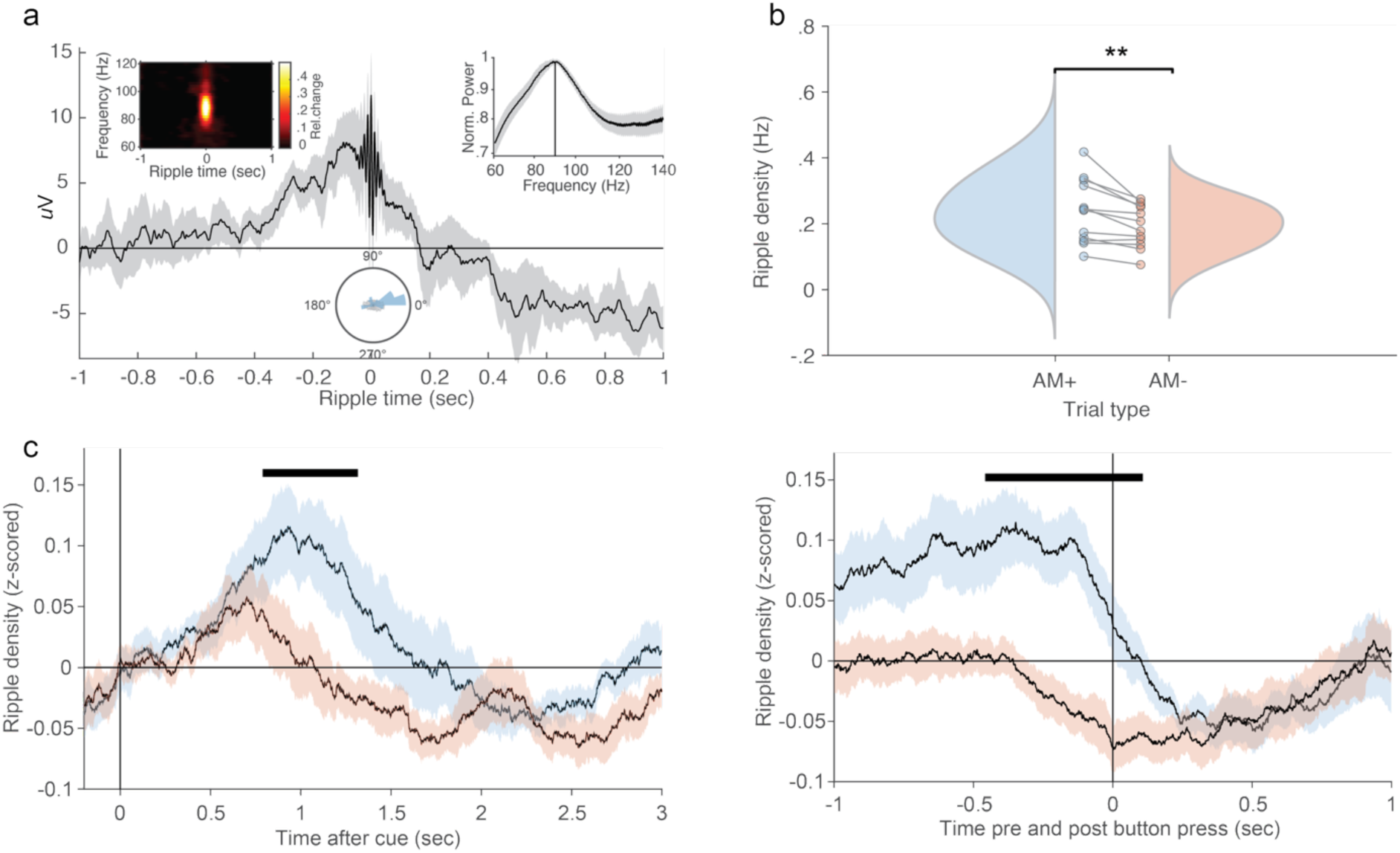
Hippocampal ripple density increases during successful memory retrieval. **a** Grand average hippocampal ripple for all participants, aligned to ripple peak at time 0. Inset: Power spectrum on ripples aligned data (left) and 1/f-corrected spectral grand average on stimulus-locked data (0-3 seconds during retrieval; right), showing peaks at 89 Hz and 87 Hz, respectively, as well as ripple peak locked to delta phase, with a mean angle of 37 degrees. **b** Ripple density significantly differed between AM+ and AM-. Circles represent participants. **c** Ripple density dynamics across time. ***Left***: aligned to retrieval cue onset. ***Right***: aligned to response time. Horizontal black lines denote significant differences between AM+ and AM- (*p* < .05, corrected for multiple comparisons across time).

Ripple density increases prior to successful free recall performance (e.g., ^33,35^). To assess whether this finding extends to associative recognition as employed here, we extracted the ripple density for each trial, i.e., the number of ripples normalised by the trial’s reaction time, and averaged across trials in each condition. Ripple density was higher during AM+ (.24 Hz, ± .03 Hz, mean ± SE) compared to AM- (.19 Hz, ± .02 Hz), *t*(11) = 3.58, *p* < .01 (Fig. 2b). Controlling for reaction time did not eliminate the effect: a mixed-effects model predicting raw ripple count confirmed that memory success remained a significant predictor after accounting for trial-level RT (linear LME: *t* = 4.79, *p* < .01; Poisson GLME: *t* = 6.65, *p* = < .01). An additional RT-matched subsampling analysis also yielded a significant effect (*t*(11) = 2.87, *p* = .02). Ripple density did not differ between recognised (AM+ and AM- trials collapsed) and unrecognised items (misses) (*t*(11) = 2.08, *p* = .06), indicating that ripple occurrence relates more to successful associative retrieval rather than recognition success alone.

Additionally, we examined whether the increased ripple rate occurred at specific time points during the successful retrieval and observed that ripple density was increased 800-1400ms post cue (for cue-aligned data; Fig. 2c, left) and leading up to the retrieval response (for response aligned data; Fig. 2c, right). These findings are consistent with previously reported latencies of episodic memory processes ^58^.

During retrieval, 21% of trials contained no ripples, 23% contained exactly one ripple, and 56% contained multiple ripples. Because more than half of all trials featured more than one event, we aligned all analyses to the ripple with the largest envelope (quantified as the sum of the root mean square of the ripple signal). This approach maximised temporal precision and avoided overlapping analysis windows that would otherwise introduce non-independent samples. Importantly, selecting a single ripple per trial did not bias the comparison between memory conditions: ripple size (*t*(11) = .69, *p* = .50), ripple duration (*t*(11) = .65, *p* = .53), and ripple density (*t*(11) = .73, *p* = .48) did not differ between AM+ and AM- trials.

### Cortical reinstatement is linked to hippocampal ripple events

Building on these findings, we next examined the relationship between hippocampal ripple events and the reinstatement of information from encoding. Specifically, we assessed whether cortical patterns present during encoding re-emerged around ripple events during retrieval.

To do so, for each participant, a Linear Discriminant Analysis (LDA ^59^) was trained on each time point around stimulus onset at encoding (−500 to 3000ms) and tested on each time point around ripples at retrieval (−1000 to 1000ms), using the preprocessed iEEG data. Importantly, hippocampal channels were used to detect ripples, but only extra-hippocampal channels (total = 647, 53.9 ± 6.24, mean ± SE, range = 24-93) were used to conduct the multivariate pattern analysis. This yielded a time-generalisation matrix (TGM; Figure 3a), showing reinstatement of encoding-related brain patterns during retrieval.

**Fig. 3.**
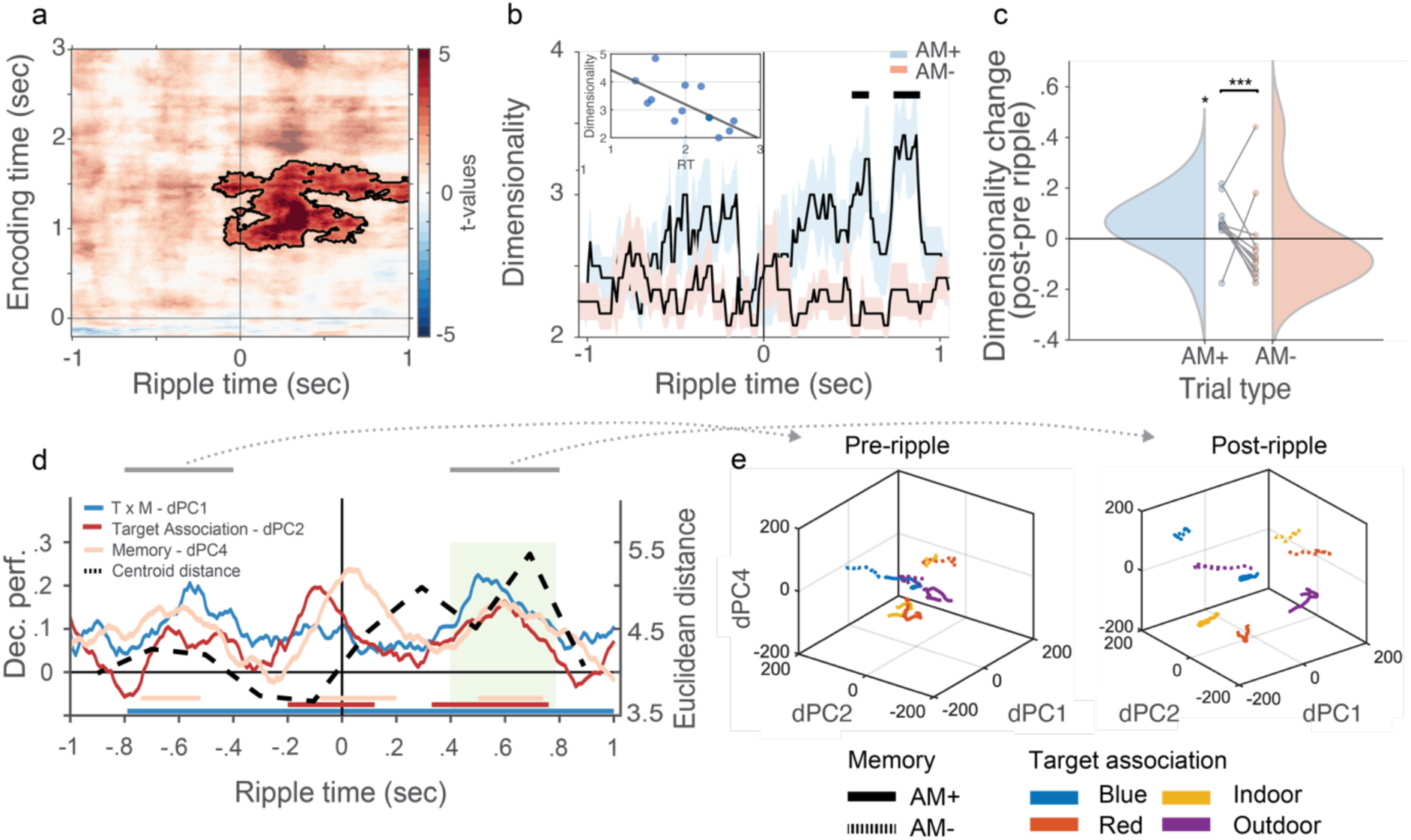
Target memory decoding and dimensionality transformation are locked to ripple events. **a** Stimulus-locked encoding x ripple-locked retrieval target decoding t-values. Results show significant increase of cortical target decodability for successful vs. unsuccessful retrieval following hippocampal ripple events (*p* < .05, corrected for multiple comparisons). **b** Estimating dimensionality for cortical contacts yielded significantly higher dimensionality for AM+ (blue) versus AM- (red) following ripple events (time 0 on x-axis). Inset: The increase in dimensionality was significantly negatively correlated with reaction time. **c** Linear mixed-effects modelling (Epoch Half: pre and post ripple, Trial Type: AM+ and AM-) of cortical dimensionality revealed a significant interaction, indicating a greater increase in dimensionality post-ripple for AM+. Asterisks denote significance compared to 0 for AM+ and the interaction of pre and post ripple time-window and AM+ and AM-. **d** dPC1, 2, and 4 revealed significant decoding accuracy of target identity in the time window of dimensionality expansion (highlighted in green bar). Note that to be able to show all components in one plot, each component’s decoding accuracy is shown as empirical data minus baseline. For each component’s decoding accuracy, including baseline, see Supplementary Fig. 9. Significant intervals of decodability are highlighted with horizontal lines. Euclidean distance between clusters increased around ripple onset with a peak 700ms post ripple-events (dashed line). **e** Plotting dPCs 1, 2 and 4, showed overlapping representations of experimental variables before ripples **(left)**, whereas their separability increased after ripples **(right)**. Note, cross-marginalisation orthogonality is not assumed in dPCA and the axes in Figure 3E represent independent sources of variance and not necessarily orthogonal subspaces.

Our results confirm that ripple-locked memory reinstatement is associated with successful associative recognition memory (AM+; p_cluster_ < .05, see Supplementary Fig. 3 for AM+ and AM-, separately, as well as when data were aligned based on cue-onset; Supplementary Fig. 4c-d for reinstatement results when dividing AM- into incorrect and don’t know responses; Supplementary Fig. 5d-e for when extra-hippocampal contact were divided into anterior and posterior regions; and Supplementary Fig. 6a for spike-corrected reinstatement). The successful decoding of memory content was closely yoked to ripple peaks in our data and continued until the end of the analysis epoch (see Supplementary Fig. 7 for various control analysis confirming that reinstatement was specific to the ripple latency of the current trial). These results extend previous work showing ripple-related reinstatement during free recall of verbal material ^35^ and suggest a broader role for hippocampal ripples’ relationship to the reinstatement of task-relevant cortical representations during associative memory retrieval.

### Hippocampal ripple-induced dimensionality expansion increases the separability of cortical representations

These results suggest that hippocampal ripples are tightly coupled to the reactivation of encoding-related cortical patterns. We propose that this ripple-locked reinstatement reflects a cortical “decompression” of memory representations, where compressed traces are expanded into distributed, high-dimensional states that support the reconstruction of latent mnemonic information during retrieval ^5,10,20,21^ (Fig. 1c). This mechanism aligns with the idea that ripples initiate a transformation in cortical dynamics, enabling the unfolding of representational content stored in compressed form. To test this, we first examined changes in representational geometry in all included extra-hippocampal channels using principal component analysis (PCA), applied to data time-locked to ripple events. More specifically, to track representational complexity over time, we applied PCA to ripple-aligned data using 60 ms sliding windows (90% overlap, −1 to +1 s). Dimensionality was estimated per window by identifying the “elbow” in the eigenvalue spectrum (second-derivative method), avoiding arbitrary variance thresholds ^60^. The number of components for each time window served as our estimate of dimensionality.

Using non-parametric cluster-based statistics, we found that cortical dimensionality significantly increased for AM+ vs. AM- trials, mainly following ripple events, with effects emerging 470-840 ms post-ripple (two clusters; both *p*<.05; see Supplementary Fig. 4a-b for dimensionality results when dividing AM- into incorrect and don’t know responses; Supplementary Fig. 5, a and b, for when extra-hippocampal contact were divided into anterior and posterior regions; Supplementary Fig. 6b for spike-corrected dimensionality). Dimensionality estimates were robust across parameters (Supplementary Fig. 8, a and b), strongly correlated with the original signal (*r_spearman_* .87, p < .001; Supplementary Fig. 8c), and AM+ trials had significantly higher dimensionality as compared to a pre-stimulus baseline (*t*(11) = 3.80, *p* = .003). Linear mixed-effects modelling confirmed higher dimensionality for AM+ trials overall (Estimate = .09, SE = .03, *t*(19,988) = 2.82, *p* = .005), and for post- vs. pre-ripple epochs (Estimate = .14, SE = .03, *t*(19,988) = 4.39, *p* < .001), with a significant interaction indicating greater ripple-induced expansion for remembered events (Estimate = -.09, SE = .02, *t*(19,988) = −4.16, *p* < .001; Fig. 3c; see Supplementary Fig. 6e-f for alternative methods to estimating dimensionality). Paired samples t-tests supported this effect, showing increased dimensionality following ripples in AM+ trials (before: 2.43 ± .05, mean ± SE; after: 2.50 ± .06; *t*(11) = 2.22, *p* = .049), but no change in AM− trials (before: 2.44 ± .06; after: 2.41 ± .06;), *t*(11) = −0.45, *p* = .66). We did not find any difference between AM+ and AM- trials in hippocampal contacts (Supplementary Fig. 9), in line with our predictions that the expansion of dimensionality takes place in cortical networks.

Crucially, higher post-ripple dimensionality during AM+ trials was associated with faster response times (*r_spearman_* = -.669, *p* = .02; Inset Fig. 3b), and stronger reinstatement (*r_spearman_* = .656, *p* = .02), linking representational expansion to both behavioural and neural markers of successful retrieval. Control analyses confirmed that these effects were specific to ripple events and not observed in surrogate data (Supplementary Fig. 10), reinforcing the functional role of ripple-triggered dimensionality expansion.

While PCA preserves the geometry of neural activity while maximising variance, and LDA enhances interpretability by maximising class separation, each has limitations. PCA mixes task-related variance and does not isolate the contribution of experimental variables, whereas LDA distorts the original geometry of the neural state space in service of classification. Demixed PCA (dPCA; ^61^), addresses both limitations by simultaneously reducing population activity and disentangling it into components aligned with specific task parameters. As illustrated in Kobak et al. ^61^, dPCA separates latent structure in a way that retains both class-specific information and the original representational geometry, making it particularly well-suited for characterising the organisation and temporal evolution of memory-related cortical states. Using this approach, we could examine how ripple events affected the organisation of cortical states, not just their dimensionality ^61^.

We reconstructed neural activity, from −1 to 1 second around ripple events, using components reflecting target association (blue, red, indoor, outdoor), memory performance (AM+ and AM-), their interaction (target association x memory performance), and a condition-independent component reflecting time. According to our hypothesis, cortical representations of these variables would separate during the period of dimensionality expansion (∼400-800 ms post-ripple; highlighted in green in Fig. 3d), thereby enhancing decodability in high-dimensional space.

We first confirmed that our reconstruction accurately captured the data. Specifically, the first 50 dPCs accounted for as much variance as our estimated total signal (see Supplementary Materials, and Supplementary Fig. 11a). To assess whether these demixed components carried task-relevant information, we applied cross-validated decoding, which tests whether the structure revealed by dPCA generalises to unseen data, providing a direct readout of when and how experimental variables are represented in neural population activity ^61^.

Each dPC is assigned to the marginalisation (target association, memory, interaction, or condition-independent) whose decoder captures the greatest portion of its explained variance. Thus, the labels in Supplementary Fig. 11b (“target association,” “memory,” etc.) directly reflect the output of the dPCA algorithm and correspond to the marginalisation that best explains each component’s variance.

Among all components, the interaction between target association and memory performance (dPC1) explained most variance (Supplementary Fig. 11b), capturing retrieval-related dynamics. This component showed high decoding accuracy across almost the entire epoch and peaked shortly after ripple events. The second component (dPC2), which captured target association-specific brain dynamics, showed significant decoding accuracy both around ripple events and during dimensionality expansion (Fig. 3d and Supplementary Fig. 11d). Finally, the fourth component (dPC4), linked to memory performance, significantly distinguished AM+ from AM- trials, with effects emerging shortly after ripple events and during dimensionality expansion (Fig. 3d and Supplementary Fig. 11f). For full output, see Supplementary Fig. 11.

These findings confirm that dPCA effectively disentangled experimental variables in a lower-dimensional space. To understand the reorganisation of cortical state space, we compared the distinctiveness of task-defined representations in pre- and post-ripple windows (−800 to - 400 ms vs. 400 to 800 ms). Using the experimentally defined labels (4 target associations × 2 memory outcomes), we computed silhouette values, where higher scores indicate stronger clustering, directly from the dPCA state space (dPCs 1, 2, and 4). This revealed a substantial increase in representational distinctiveness following ripples (pre-ripple: .632; post-ripple: .903; t(327) = 13.03, p < .01), indicating that cortical activity patterns corresponding to the eight task conditions occupy more distinct regions of state space after ripple onset (Fig. 3d).

When analysing AM+ and AM- trials separately both trial types showed a significant pre- to post-ripple increase (AM+: pre = .626, post = 0.915, t(163) = 11.39, p < .01; AM-: pre = .644, post = .892, t(163) = 7.44, p < .01), but the effect was significantly larger for correct retrieval (AM+ difference = .289; AM- difference = .249; t(163) = 2.56, p = .01), consistent with the interpretation that ripple-triggered state-space expansion supports successful memory retrieval. We observed the same pattern when analysing the alternative component set (dPCs 1, 3, and 4; Supplementary Fig. 11h). Using label-defined clusters, silhouette values again showed a marked increase from the pre- to post-ripple window (pre: .762; post: .914; t(327) = −8.25, p < .001). Both AM+ and AM- trials exhibited significant increases in separability (AM+: pre = .748, post = .925, t(163) = 6.73, p < .001; AM-: pre = .776, post = .904, t(163) = 4.95, p < .001). Importantly, the ripple-related increase in separability was significantly larger for remembered than forgotten trials (AM+ difference = .177; AM- difference = .128: t(163) = 6.64, p < .001).

These findings confirm that dPCA effectively disentangled experimental variables in a lower-dimensional space and that ripple onset is followed by a robust reorganisation of cortical state space, with stronger distinctiveness of task-relevant structure during successful memory retrieval.

To quantify this representational reorganisation, we applied k-means clustering to the dPCA components and assessed the Euclidean distance between clusters. To evaluate clustering quality, we used the Silhouette score. Using 200 ms time bins around ripple events, we observed increased cluster separation beginning at ripple onset and peaking ∼700 ms later (Fig. 3d, dashed line). To further quantify this effect, we plotted the state-space organisation of these components during dimensionality expansion in the same time windows as in the previous analysis (400-800ms pre- and post-ripple events). Before ripple events, we identified six clusters (silhouette score = .83), whereas eight clusters emerged after ripple events (silhouette score = .90) (Fig. 3e). This was accompanied by a significant increase in Euclidean distance between clusters (pre-ripple: 4.06 ± .044, mean ± SE; post-ripple: 4.78 ± .048; t(319) = 5.04, p < .01). A similar pattern emerged when using dPC 1, 3, and 4. We observed that the optimal cluster count increased from four clusters pre-ripple (silhouette score = .85) to eight clusters post-ripple (silhouette score = 0.91). The Euclidean distance between cluster centroids also significantly increased (pre-ripple: 3.51 ± .043; post-ripple: 4.66 ± .052; t(319) = 7.12, p < .01; Supplementary Fig. 11h). In both visualisations, we observed a clear separation of all experimental variables following ripple events, a pattern that was less distinct before ripple onset.

### Theta-gamma phase-amplitude coupling coordinates information flow between hippocampus and cortex

Our results demonstrate that hippocampal ripples are associated with a cortical expansion during episodic memory retrieval in which the separation between experimental variables increases. But what mechanisms enable this transformation of information? A candidate mechanism is the coupling between hippocampal theta and cortical gamma rhythms ^40,41,46,62–64^. Despite theoretical accounts emphasising its importance, there is currently little direct evidence in humans that theta-gamma-phase-amplitude-coupling (TG-PAC) supports statistical dependencies between regions during episodic memory retrieval ^65^.

One likely reason for this gap is the considerable variability in retrieval timing across trials, which poses a challenge for aligning neural events consistently. Our findings suggest that hippocampal ripples can be leveraged to pinpoint the exact moments of reinstatement. Thus, we hypothesised that any statistical marker of coordinated activity between hippocampus and cortex would occur shortly after ripple onset, but prior to the cortical expansion of memory representations, and would be critical for reconstructing the memory trace. To test this, we examined TG-PAC in the time window surrounding ripple events.

We identified peak theta in the hippocampal signal and gamma frequencies in the extra-hippocampal signal after removing the 1/f background using the FOOOF algorithm ^62^. Time-frequency data were then aligned to these peaks (±2 Hz for theta, ±10 Hz for gamma), and phase-amplitude coupling was computed per channel pair which was then averaged. Empirical PAC values were compared to a null distribution generated by shuffling trials 500 times per participant and time point. This allowed us to assess TG-PAC in narrow-band activity centred on peak frequencies, time-locked to ripple events during retrieval (−1 to +1 s). In addition, we quantified gamma waveform-shape metrics (peak-trough sharpness, rise-decay asymmetry, skewness, kurtosis) across regions and gamma sub-bands. These analyses confirmed that gamma waveforms were highly symmetric and did not vary systematically across cortical regions or frequencies, making it unlikely that the observed TG-PAC effects were driven by waveform shape rather than genuine hippocampal-cortical coupling (Supplementary Fig. 12).

The average peak theta and gamma frequencies were 6.08 Hz (± .69 Hz) and 59.58 Hz (± 4.75 Hz), respectively (Fig. 4a). To assess the appropriateness of using participant-level peak frequencies for PAC alignment, we quantified the dispersion of per-channel peaks around averaged theta and gamma peaks. Hippocampal theta peaks showed very low variability, whereas cortical gamma peaks showed broader dispersion, as expected for distributed cortical sites ^66^, though the majority of channels lay close to the participant-level peak and only a small subset of high-frequency outliers accounted for the wider spread (See Methods for details).

**Fig. 4.**
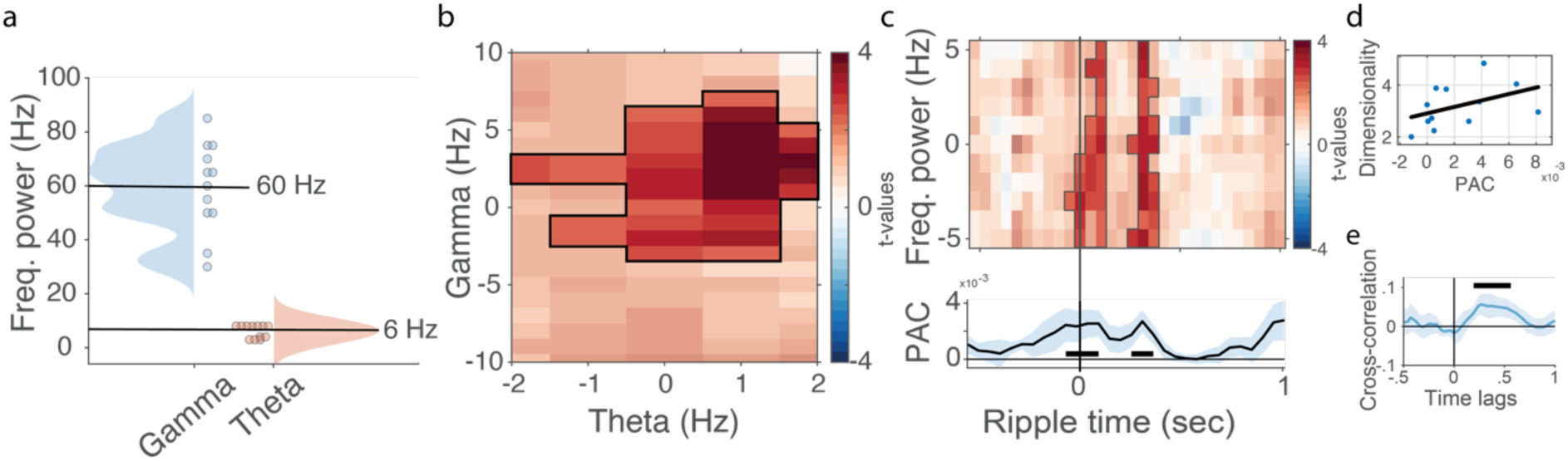
Phase-amplitude coupling following ripple events are related to dimensionality expansion. **a** On average gamma oscillations peaked at 60 Hz, whereas theta on average peaked at 6 Hz. **b** Significant TG-PAC around peak frequencies for respective frequency (0 on x and y axes). **c**. Top: A time-resolved TG-PAC commencing around ripple events and persisted for approximately 400ms after the events. Bottom: Plotting the time-resolved TG-PAC averaged across the y-axis in top panel. **d** A positive correlation between TG-PAC and cortical dimensionality expansion, such that stronger TG-PAC was related increased dimensionality expansion. **e** A cross-correlation showed that TG-PAC leads dimensionality expansion with a peak at approximately 260ms.

We observed a significant coupling between hippocampal theta phase and cortical gamma amplitude around the identified peaks when contrasting AM+ with the shuffled control (*p_cluster_*<.05; Fig. 4b). We did not find a significant difference between AM+ and AM- trials, suggesting that TG-PAC is as a general feature of retrieval-related hippocampal-cortical coordination rather than a memory-specific effect.

To examine the timing of TG-PAC relative to ripple events, we conducted a time-resolved analysis using sliding windows (same settings as in the previous analysis; see Methods for details). This revealed that coupling emerged shortly after ripple onset and persisted for ∼300 ms (*p_clusters_* < .05; Fig. 4c). For each participant, we averaged over the significant time points for TG-PAC and dimensionality, separately (black horizontal lines), and correlated the resulting vectors. This analysis showed that TG-PAC strength in this post-ripple window correlated with ripple-related increases in cortical dimensionality (r = .59, p = .04; Fig. 4d), indicating an association between cross-frequency coupling and representational transformation during retrieval.

We hypothesised that TG-PAC would follow hippocampal ripples but precede cortical dimensionality expansion. To test this, we performed a cross-correlation analysis against a null distribution of 1000 phase-shuffled surrogates. The analysis revealed a significant positive lag, peaking at ∼260 ms (Fig. 4e), suggesting that TG-PAC may reflect an intermediate process occurring between hippocampal output and cortical reorganisation.

## Discussion

Episodic memory is thought to optimise neural processing by compressing information during encoding and expanding it during retrieval ^3–6^. The hippocampus plays a central role in this dynamic, matching retrieval cues with stored memory traces and triggering pattern completion to reconstruct past experience in the cortex ^9–11^. Yet, the neural mechanisms by which hippocampal output enables this transformation remain poorly understood.

We hypothesised that memory retrieval involves a ripple-related transformation in neural dimensionality, from low-dimensional representations in the hippocampus to high-dimensional cortical activity patterns. Such a shift would increase representational capacity, allow cortex to differentiate features, minimise interference, and flexibly decode mnemonic content ^5,10,20,21^.

To test this, we analysed human intracranial EEG (iEEG) data from 12 participants performing an associative recognition memory task. We found that that successful memory retrieval was associated with stronger reactivation of encoding-related cortical patterns (Fig. 3a), and crucially, with increased cortical dimensionality (Fig. 3, b-c). This expansion was linked to both faster response times (Fig. 3b) and greater reinstatement strength, suggesting a functional role for dimensional expansion in supporting retrieval.

Our findings position hippocampal ripples as key triggers of cortical reinstatement during episodic memory retrieval. While prior work has linked ripples to hippocampal-cortical communication and memory-related activity in both humans and non-human animals ^30,31,33–36,67–70^, we show that ripples are temporally aligned with both increased cortical dimensionality and reinstatement of encoded information. This extends previous literature by providing direct evidence that ripples are associated with transition from compressed to high-dimensional cortical states supporting the reconstruction of specific mnemonic content. Moreover, our results suggest that ripple-related reflect a broader cortical mechanism for memory retrieval.

A potential concern is that interictal epileptiform discharges (IEDs) themselves can drive hippocampus-cortex coupling, as demonstrated by Gelinas et al. 2016 ^71^. This raises the possibility that ripple-related effects could, in principle, reflect IED-related dynamics rather than physiological ripple activity. However, several aspects of our data argue against this interpretation. First, ripple detection was restricted to hippocampal channels, whereas cortical dimensionality and reinstatement were assessed in anatomically distinct, non-hippocampal regions, reducing the likelihood that local epileptiform activity directly drives the observed cortical effects. Second, we explicitly controlled for interictal activity by excluding 1 second of data surrounding detected spikes across all channels, and the results remained unchanged (Supplementary Fig. 6). Third, when conducting the main analyses with epochs realigned to identified spikes, we did not find any difference between conditions (Supplementary Fig. 15). Fourth, interictal spikes are typically brief, spatially heterogeneous events that do not induce sustained, globally synchronous activity across distributed cortical regions. In contrast, the ripple-related effects we observe unfold over several hundred milliseconds and are associated with structured, condition-specific changes in cortical representational geometry. Together, these considerations suggest that the reported hippocampal-cortical interactions are unlikely to be explained by IED-related coupling and instead reflect physiological ripple-mediated communication.

We identified hippocampal ripples to be related to this cortical expansion. Ripple events were more frequent on successful trials and reliably preceded both reinstatement and increased dimensionality. This finding supports a novel account of ripple events: rather than merely reactivating previously stored patterns, ripples appear to also be associated to a transformation that reconstructs compressed memory representations into complex, high-dimensional cortical states. Such expansion may reflect the recruitment of a broader set of cortical areas and the unfolding of more differentiated neural activity patterns, enabling flexible decoding of specific mnemonic content. This interpretation is consistent with extensive rodent literature showing that hippocampal ripples carry time-compressed replay of recent experience ^72–74^. Such compressed ripple content provides an efficient substrate for broadcasting mnemonic information to cortex, where it can be expanded into high-dimensional representations that support detailed reconstruction.

Recent studies have examined the temporal relationship between hippocampal activity and cortical reinstatement. Michelmann and colleagues ^75,76^ showed that when analyses are anchored to cortical reinstatement events in sensory regions, hippocampal activity reliably precedes cortical reinstatement by approximately 500–740 ms during memory retrieval. Importantly, these studies do not align reinstatement to internally generated hippocampal output events themselves. In contrast, our analysis is explicitly ripple-locked. When aligned to hippocampal ripple peaks, we observe reinstatement emerging immediately following the ripple, followed by a broader increase in cortical dimensionality that peaks approximately 400–800 ms later, and this effect was stronger in posterior as compared to anterior channels (Supplementary Fig. 5d-e), in line with recent work on transformation of memory traces during episodic memory retrieval (Rau et al., 2025). Thus, our findings complement those of Michelmann et al. by showing that reinstatement begins rapidly following hippocampal ripples, while large-scale cortical reorganisation unfolds over several hundred milliseconds, consistent with multi-synaptic propagation across cortical networks ^77,78^.

To characterise this reorganisation, we used demixed PCA (dPCA). Task-relevant variables (e.g., memory accuracy and target identity) became more separable following ripples (Figure 3e), with maximal separation aligning temporally with the rise in dimensionality (Figure 3d). The neural state space also became more complex, as indicated by more optimal clusters and greater distances between them. This supports the idea of ripple-related representational "decompression".

Additionally, when analysing the distinctiveness of task-defined representations, we observed a systematic increase in silhouette values following ripples, indicating that cortical activity patterns associated with the eight task conditions occupied more well-defined and segregated regions of state space. Crucially, both AM+ and AM- trials showed significant increases in distinctiveness, but AM+ trials exhibited a reliably larger ripple-related improvement. This pattern indicates that ripple onset is linked to a general reorganisation of cortical geometry, where successful retrieval is associated with a stronger, more selective distinctiveness of task-relevant structure.

We also report the first evidence of hippocampal theta-cortical gamma phase-amplitude coupling (TG-PAC) during episodic memory retrieval in humans. TG-PAC peaked shortly after ripple onset (Figure 4c) and preceded cortical dimensionality expansion by ∼260 ms (Figure 4e), suggesting it may serve as a bridge between hippocampal output and cortical reorganisation. This builds on work showing TG-PAC during hippocampal encoding/retrieval ^47,49,64,79^, spatial memory ^80^, and autobiographical recall ^81^.

While some studies have linked slow vs. fast gamma to encoding and retrieval respectively ^46,82,83^, we found no consistent frequency separation across participants. This may reflect region-specific gamma profiles outside the medial temporal lobe. Importantly, we also acknowledge that our cortical gamma-band likely includes contributions from high-frequency events in the ripple range; rather than assuming strict separation between gamma and cortical ripples, we interpret our findings as evidence that hippocampal theta rhythm gates a broad class of high-frequency cortical activity, including potential ripples, that precedes and predicts ripple-locked increases in cortical dimensionality ^35^.

Our results contribute to a broader literature on neural dimensionality as a mechanism for cognitive flexibility. Studies in decision-making and navigation have shown that high-dimensional states support adaptive coding (e.g., ^13^). We extend these findings to episodic memory, showing that successful retrieval engages dynamic, high-dimensional cortical representations ^5^. Prior work has also shown that events with greater encoding dimensionality are more likely to be remembered ^84^, raising the possibility that rich encoding geometry facilitates later reconstruction. Future work should test whether dimensionality at encoding predicts reinstatement fidelity.

We propose a framework in which ripples trigger a shift from compressed hippocampal codes to expanded cortical states. This transition may be mediated by ripple-induced synchrony ^85^, followed by cortical desynchronisation in alpha/beta bands ^86^, which has been linked to improved memory fidelity and recall ^80,81^. Such desynchronisation may support low-synchrony, high-dimensional representations that promote selective decoding while reducing interference _87–91._

Understanding how ripple-induced synchrony interacts with cortical desynchronisation will be an important direction for future research on the temporal dynamics of memory reinstatement.

## Acknowledgement

The authors would like to thank Juergen Fell and Bernhard Staresina for sharing their data.

## Funding

Max Planck Society (C.K. and C.F.D.)

Deutsche Forschungsgemeinschaft project 437219953 (S.M.)

## Author contributions

Conceptualization: C.K.

Methodology: C.K. and S.M.

Investigation: C.K.

Visualization: C.K.

Funding acquisition: S.M. and C.F.D.

Project administration: C.F.D.

Supervision: S.M. and C.F.D.

Writing – original draft: C.K., S.M. and C.F.D.

Writing – review & editing: C.K., S.M. and C.F.D.

## Declaration of interests

The authors declare no competing interests.

## Resource availability

### Lead contact

Requests for further information should be directed to and will be fulfilled by the lead contact, Casper Kerrén.

### Materials availability

The study did not generate new, unique reagents.

### Data and code availability

The data and code that support the conclusions of this study are available online at GitHub (code: https://github.com/kerrencasper/Hippocampal-ripples-initiate-cortical-dimensionality-expansion-for-memory-retrieval.git) and Zenodo (data: 10.5281/zenodo.18490239).

## Methods

### Experimental method and study participant details

#### Participants

A total of 15 patients took part in the study. Three were excluded from further analysis due to clinical monitoring revealing epileptogenic activity in both hippocampi, resulting in a final sample of 12 patients (6 female; 33 years ± 9.3, mean ± SD) with pharmacoresistant epilepsy. All participants provided written informed consent, and the study was approved by the Ethics Committee of the Medical Faculty at the University of Bonn. Data were recorded at the Department of Epileptology, University Hospital Bonn. Complementary analyses from a subset of participants in the current paradigm have been reported previously: 5 patients in an earlier study ^92^ and 11 patients in a later study ^93^.

### Method details

#### Experimental procedures

Participants were seated upright in a sound-attenuated room, approximately 50 cm from a laptop screen, and engaged in an associative learning paradigm (Fig. 1a). Each experimental block consisted of an encoding phase, a 1-minute distractor phase, and a retrieval phase.

During encoding, participants were presented with a German noun paired with either a colour or a scene, depending on the run. Colour and scene runs alternated: in colour runs, the noun was paired with either a red or blue square; in scene runs, it was paired with an image depicting either an indoor (e.g. an office) or outdoor (e.g. a nature scene) environment. The task was to form an association between the word and the accompanying stimulus by vividly imagining the object described by the noun in conjunction with the colour or scene (e.g. “a red lemon” or “an elephant in the mountains”), and to rate the plausibility of the imagined scenario. This means that multiple words were associated with the same picture. However, forming individual association. Participants had up to 3 seconds to make their plausibility judgement via button press. Each trial was preceded by a jittered inter-trial interval (ITI) of 700-1300 ms (mean = 1000 ms), during which a fixation cross was displayed at the centre of the screen. Trials ended immediately upon response.

In the retrieval phase, participants were presented with 50 previously seen words randomly intermixed with 25 novel words, along with four response options. Their task was to indicate whether the word was new (‘N’ response), whether it was old but the associated target could not be recalled (‘?’ response), or whether it was old and the associated colour or scene could be correctly retrieved (in which case the appropriate response option was selected).

Responses were self-paced with an upper time limit of 5 seconds. As in encoding, trials were terminated by a button press and were preceded and followed by a jittered ITI (700-1300 ms, mean = 1000 ms) showing a fixation cross. Each run lasted approximately 9 minutes.

#### Implantation of depth electrodes

Intracranial electroencephalography (iEEG) data were referenced to linked mastoids and recorded from medial temporal lobe regions, including the hippocampus, as well as additional cortical areas, at a sampling rate of 1 kHz (bandpass filter: 0.01 Hz to 300 Hz) (Fig. 1b). Depth electrodes targeting the hippocampus were implanted stereotactically as part of presurgical evaluation, following two different implantation schemes: in eight participants, electrodes were placed along the longitudinal axis of the hippocampus, while in four participants, electrodes were implanted laterally via the temporal lobe. All participants were on anticonvulsive medication, with plasma levels maintained within the therapeutic range. In addition to depth electrodes, scalp electrodes were placed at positions Cz, C3, C4, and Oz according to the international 10-20 system; however, these were excluded from all subsequent analyses.

#### Electrode selection

Electrode contact localisation was determined using multiple complementary criteria. First, we inspected post-implantation MRI scans and identified electrode contacts located within the hippocampus. Second, pairwise channel coherence in the 4-8 Hz range was calculated during the retrieval phase, based on the assumption that contacts within the same anatomical region would exhibit high coherence ^94,95^. Third, event-related potentials (ERPs) were computed for each contact, with the expectation that electrodes in the same region would display similar ERP profiles.

Only hippocampal contacts from the clinically defined healthy hemisphere were included (for one participant, both hemispheres were considered healthy). Following preprocessing (see below) and the application of these localisation criteria, a total of 72 hippocampal contacts across 12 participants were identified as clean and reliably located (6.0 ± .64, mean ± SE contacts per participant). For extra-hippocampal regions, 647 contacts were retained for analysis (53.9 ± 6.24 per participant). See Fig. 1b for a summary across participants and Supplementary Fig. 1 and 2 for individual electrode maps.

In cortical regions, across participants, the highest coverage was observed in left inferior temporal gyrus (AAL atlas: Temporal_Inf_L; 84 electrodes in 12/12 patients), right parahippocampal gyrus (ParaHippocampal_R; 78 electrodes in 12/12 patients), left parahippocampal gyrus (ParaHippocampal_L; 76 electrodes in 12/12 patients), and right inferior temporal gyrus (Temporal_Inf_R; 76 electrodes in 11/12 patients). These regions are strongly implicated in visual and associative/semantic memory processing.

#### Preprocessing

Data processing was carried out using FieldTrip (version 20230422; ^90^) standard MATLAB functions, and custom-written MATLAB scripts. Line noise was removed using 2-Hz-wide bandstop filters centred at 50, 100, 150, and 200 Hz. Following this, the data were re-referenced using a common trimmed average approach, implemented via MATLAB’s *trimmean* function. Specifically, 20% of the highest and lowest values were trimmed to reduce the influence of outliers before computing the mean, which was then subtracted from each channel.

We selected this method of referencing because dimensionality-based multivariate analyses (PCA, dPCA) rely on preserving shared variance across spatially distributed signals, and trimmed averaging provides a robust estimate of the global reference while avoiding artefact-driven bias. Importantly, we verified that all key ripple characteristics and multivariate results remained stable using bipolar referencing (Supplementary Fig. 13).

We applied an automated artefact rejection procedure adapted from standard intracranial EEG preprocessing protocols ^35,96^. For each channel, the continuous signal was z-scored according to three metrics: (i) absolute amplitude, (ii) the sample-to-sample voltage gradient, and (iii) high-frequency amplitude obtained after high-pass filtering at 250 Hz to detect epileptiform spikes. A data point was labelled as artefactual if it exceeded a z-score of 6 on any single metric or a z-score of 4 on a conjunction of two metrics (gradient + high-frequency, or absolute amplitude + high-frequency). To ensure conservative rejection, we removed an additional 50 ms of data on either side of each marked segment before any analyses.

#### Ripple detection

Ripple events were detected independently in each hippocampal channel using a validated ripple detection algorithm ^96^. Channel-specific artefact segments, including epileptogenic spikes detected by an automated algorithm, were excluded from the analysis with an additional ±50 ms padding window. Only data from non-pathological hemispheres were analysed.

For each channel, the signal was bandpass filtered between 80 and 120 Hz using a two-pass FIR filter. A 20 ms root-mean-square (RMS) envelope of the band-passed signal was then computed and smoothed with a 20 ms moving window. Ripple detection thresholds were computed in a strictly channel-wise fashion using only artefact-free data: the main detection threshold was set to the mean ripple-band envelope plus 1.5 standard deviations, and an upper cutoff of mean + 9 standard deviations was applied to suppress large-amplitude transients during threshold estimation.

Continuous supra-threshold segments were then identified and retained only if they met standard physiological criteria. Events had to be between 38 and 500 ms in duration and had to contain at least three full oscillatory cycles (three peaks and three troughs) in the underlying raw or band-passed signal, ensuring that events reflected true high-frequency bursts rather than single sharp deflections. For additional confirmation, each candidate event underwent a false-positive rejection procedure based on its frequency profile: for every event, a time-frequency representation (65-135 Hz) was computed in a ±250 ms window around the envelope peak, and the spectral power was normalised across frequencies. Events were accepted only if they exhibited a prominent narrowband peak within 75-125 Hz, providing independent verification that the event was spectrally consistent with physiological ripple activity. Events failing any of the criteria above were discarded.

Together, these steps reassured that we found physiological ripples, rather than pathological ones or artifacts.

Ripple density was calculated by dividing the number of detected events by the length of the corresponding trial, up to the participant’s reaction time. As ripple detection could not be performed during artefactual segments, any time marked as artefact was subtracted from the trial length prior to this calculation. The resulting value reflects the frequency of ripple occurrence per trial. Since RT during AM+ trials were slightly faster than during AM- trials, we conducted additional control analyses to ensure that ripple density was not explained by RT differences.

First, we examined raw ripple counts per trial (summing events across hippocampal channels, without any averaging or RT normalisation). AM+ trials contained numerically more ripples (M = 2.784) than AM- trials (M = 2.457).

We used trial-wise mixed-effects regression to predict raw ripple counts while statistically controlling for RT, with subject included as a random intercept. Memory condition (AM+/AM-) remained a highly significant predictor in both a Gaussian linear mixed model (estimate = .473, t = 4.793, p < .01) and a Poisson generalised linear mixed model (estimate = .195, t = 6.651, p < .01). Thus, the association between memory success and ripple occurrence persists when RT is explicitly modelled as a covariate at the level of individual trials.

To rule out any residual RT confounds, we performed an RT-matched subsampling analysis in which AM+ and AM- trials were equated in their RT distributions within each participant using quantile-based matching. Even with RT fully controlled in this way, AM+ trials continued to exhibit significantly higher ripple counts (t(11) = 2.868, p = .015).

Together, these convergent analyses demonstrate that the memory-related increase in hippocampal ripple occurrence is not an artefact of RT differences. Rather, ripple activity is genuinely enriched on successful associative memory trials, regardless of whether ripples are quantified as density, raw counts, regression-adjusted estimates, or RT-matched subsamples.

For all main analyses, we selected a single ripple per trial; the one with the maximum envelope, computed as the sum of the root mean square (RMS) of the ripple signal. During retrieval, 21% of the trials contained no ripples, 23% contained exactly one ripple, and 56% contained more than one ripple. Across trials, the median ripple occurred at 1002 ms after cue onset during retrieval.

To examine whether the increase in ripple rate was time-specific, we conducted a time-resolved ripple-rate analysis. Data were aligned to both cue onset and reaction time during retrieval. For each channel, the ripple time series was smoothed with a 400 ms moving average using MATLAB’s *smoothdata* function, and z-scored across all conditions. We then collapsed across channels and conditions and performed a cluster-based permutation test to identify significant changes in ripple rate over time.

#### Multivariate pattern analysis

The raw iEEG time series were epoched based on two time points: trial onset during encoding and ripple onset during retrieval. This yielded two separate datasets: the encoding-aligned data, used as the training set (from −500 ms to +3000 ms relative to cue onset), and the ripple-aligned data, used as the testing set (from −1000 ms to +1000 ms relative to ripple onset). Importantly, hippocampal channels were used to detect ripples, but only cortical channels were used to conduct the multivariate pattern analysis.

Encoding and ripple-aligned data were downsampled to 100 Hz, smoothed using a 200 ms moving average (using the MATLAB function *smoothdata*) and baseline-corrected using a 200 to 0 ms pre-cue interval. For ripple-aligned retrieval data, the baseline was defined using the pre-cue window from the corresponding encoding trial in which the ripple occurred.

All encoding trials were included in classifier training and split into four stimulus classes: blue vs red (for colour runs) and indoor vs outdoor (for scene runs). These classes were used to train separate classifiers. For retrieval, classification performance was assessed separately for trials with successful associative memory (AM+) and unsuccessful memory (AM-, including both incorrect and ‘don’t know’ responses). After artefact rejection, the average number of trials per participant was similar across conditions (AM+: 58.67 ± 32.05, mean ± SD; AM-: 57 ± 31.87; t(11) = 0.11, p = .91). ‘Don’t know’ trials were included for two reasons: (1) to better match the number of trials across conditions, particularly for later dimensionality analyses, and (2) because failure to recall is behaviourally equivalent to an incorrect memory judgement. Note that when dividing AM- into incorrect and ‘don’t know’ responses for Supplementary Figure 4, there was a very limited number of incorrect trials for some participants and these results should therefore be interpreted with caution.

Prior to classification, both training and testing datasets were z-scored independently. A linear discriminant analysis (LDA), implemented via the MVPA-Light toolbox ^59^, was used to train and test the classifier at each time point, yielding a time-generalisation matrix (TGM). As the training and testing sets were drawn from independent datasets (encoding vs retrieval), no cross-validation was performed. Statistical comparisons between conditions were assessed using cluster-based permutation tests.

To confirm that reinstatement was specifically linked to the timing of ripple events, we ran three control analyses involving trial-wise ripple time shuffling. First, from 1-5 seconds after cue onset during retrieval we picked 60 time points, which we treated as fictitious ripple events. We aligned the data to these events and ran the dimensionality analysis from −1 to 1 second around the events (RetrCorrect). Second, ripple times were randomly reassigned across AM+ trials 1000 times per participant (RandCorrect). Third, ripple times were circularly shifted to the next trial using MATLAB’s *circshift* function (CircCorrect), providing a stricter temporal control (Supplementary Fig. 7).

For the first two control analyses, statistical significance was assessed using a group-level permutation framework following ^97^. Empirical values were first averaged within the time-frequency cluster identified in the main analysis, yielding a single cluster-mean value per participant. A group-level null distribution was then constructed by repeatedly sampling one surrogate estimate per participant from the corresponding control distribution and averaging these values across participants over 10 000 iterations. The empirical group-mean cluster value was compared against this null distribution to obtain a two-sided p-value.

For the first control (RetrCorrect), empirical values were significantly greater than the group-level null distribution (z = 6.05, p < .01, one-sided). For the second control (RandCorrect), empirical values again exceeded the null distribution (z = 2.19, p = .029, one-sided). As no comparable group-level surrogate distribution was available for the CircCorrect control, empirical values within the significant cluster were compared directly with the circularly varied data using a paired-samples t-test. This analysis showed significant stronger reinstatement for the empirical data relative to CircCorrect (t(11) = 2.05, p = .03, one-sided).

#### Dimensionality transformation

The same procedure as described above was used to epoch and preprocess the data. All subsequent analyses were conducted separately for AM+ and AM- trials. Ripple-aligned data were segmented into temporal windows using a 60 ms sliding window with 90% overlap, spanning from −1 to +1 second around ripple onset. This windowing approach was chosen to increase the signal-to-noise ratio for dimensionality estimation and to produce a smooth temporal profile of representational complexity. Similar results were observed when using 100 ms or 200 ms sliding windows (see Supplementary Fig. 8a and b).

For each time window, we estimated the embedding dimensionality using principal component analysis (PCA). The eigenvalues of the covariance matrix were extracted, and the number of retained components was determined via the second derivative method (i.e., identifying the "elbow" point at which the explained variance sharply declined across subsequent components). This data-driven approach avoids the arbitrary selection of a fixed variance threshold (e.g., 85-90%), which is known to be problematic in PCA analyses ^60^. The number of retained components for each time window served as our estimate of dimensionality (variance explained within significant time window, AM+: 53.85% ± 10.06%, mean ± SD; AM-: 46.87% ± 13.17%; paired samples t-test between trial types: t(1,11) = 2.80, p = .02).

To ensure that our dimensionality estimates did not depend on the elbow-detection procedure, we implemented an additional control analysis based on the deviation of each eigenvalue spectrum from a fitted power-law distribution. For each time window, we fitted a power-law function to the normalised eigenvalues of the covariance matrix using least-squares regression in log-log space (MATLAB fit, model power1). We then quantified the strength of the elbow by computing the signed difference between the empirical eigenvalue at the elbow (identified via the second-derivative method) and the eigenvalue predicted by the fitted power-law at the same component index. Large positive values indicate that the empirical spectrum bends more sharply than expected under a scale-free distribution, reflecting a stronger separation between signal- and noise-dominated components, whereas values close to zero indicate an eigenvalue decay well-approximated by a power-law.

Critically, this metric does not provide an independent estimate of dimensionality; rather, it quantifies how structured or non-power-law-like the eigenvalue spectrum is at each time point. The temporal profile of this measure closely paralleled the dimensionality time course obtained using the elbow method (Supplementary Fig. 8d): both showed a marked increase following ripple events, and both were significantly larger for AM+ compared with AM- trials. This convergence demonstrates that the observed ripple-triggered increase in representational complexity is not an artefact of the elbow-detection algorithm but reflects a genuine change in the structure of the neural covariance spectrum.

To statistically assess the temporal difference in dimensionality between AM+ and AM- trials, we applied cluster-based permutation testing. Additionally, to confirm that dimensionality expansion was specifically linked to ripple timing, we repeated the same control analyses used in the reinstatement analyses (see Supplementary Fig. 10). All three control comparisons revealed significantly greater dimensionality in the empirical data compared to the shuffled conditions (RetrCorrect: t(11) = 3.97, p < .01; RandCorrect: t(11) = 3.78, p < .01; CircCorrect: t(11) = 3.36, p < .01; all one-sided).

In a follow-up analysis, per participant, we extracted the average dimensionality estimate within the significant time points (black horizontal line in Figure 3b) and correlated this resultant vector containing one number per participant with (1) participants’ reaction times (correct trials only), and (2) decoding accuracy (i.e., classifier performance for correct minus incorrect trials) using two-sided Spearman’s rank correlations.

To further assess and to gain more specificity of the dimensionality transformation, we split the data into blocks (3-6 blocks per participant) and fitted a linear mixed effect model with the fixed effects being conditions (AM+ and AM-), dimensionality and time around ripple (pre and post [-1 to 0 and 0 to 1]) (Supplementary Table 2). Random effects were participant and blocks. We used the MATLAB function *fitlme* for the model fitting.

### Control analyses

Comparable results were obtained when using effective dimensionality (ED; Supplementary Fig. 8e), an alternative metric that estimates the intrinsic dimensionality of neural population activity ^98^:

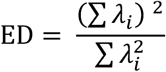

where *λ* are the eigenvalues (variance explained by each principal component).

We also estimated the number of signal-dominated components using a Marčenko-Pastur (MP) threshold ^99^, which provides a theoretically motivated upper bound on eigenvalues expected from random, uncorrelated data. For each sliding time window, we z-scored neural activity channel-wise and computed the channel × channel correlation matrix,

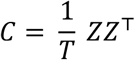

where Z is the *n*_chan_ X ⊤ matrix of z-scored activity and ⊤ is the number of time points in the window. We then obtained the eigenvalue spectrum of C and sorted eigenvalues in descending order. Under the null hypothesis of independent noise, eigenvalues of random correlation matrices follow the Marčenko-Pastur distribution with an upper bound:

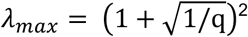

where q = ⊤ / *n*_chan_ ≥ 1. Because z-scoring sets *σ*^2^ =1, this expression gives the theoretical maximum eigenvalue expected from noise. All eigenvalues exceeding *λ*_$%&_were therefore classified as signal components, and the number of such suprathreshold eigenvalues (i.e., ∑*_i_ λ_i_*> *λ_max_*) provided a data-driven estimate of the effective rank or intrinsic dimensionality of the population activity.

This MP-derived dimensionality measure yielded patterns highly consistent with our PCA-based and ED-based analyses, confirming that ripple-locked increases in cortical dimensionality reflect genuine structure in the neural data rather than noise. Linear mixed-effects modelling revealed that cortical dimensionality was significantly higher in the post-ripple window compared to the pre-ripple window (Estimate = .17, SE = .047, t = 3.64, p < .01) Although the main effect of memory condition (AM+ vs AM-) did not reach significance (Estimate = -.089, SE = .047, t = −1.90, p = .057), there was a significant Condition × Half interaction (Estimate = -.083, SE = .030, t = −2.82, p = .0049), indicating that the post-ripple increase in dimensionality was significantly larger for remembered (AM+) than forgotten (AM-) trials (Supplementary Fig. 8f).

#### Reconstruction of original data and decoding

To ensure that dimensionality reduction preserved task-relevant structure in the neural data, we performed a reconstruction analysis that quantified how much of the original decodable information was retained after projecting the data into the reduced PCA space.

First, for each participant we computed the within- and between-class covariance matrices and extracted the eigenvectors corresponding to the number of PCA components identified through the elbow method. We then reconstructed the neural activity by multiplying the class centroids for each condition with these eigenvectors, separately for encoding and retrieval. This produced a low-dimensional approximation of the original data expressed back in the full sensor space.

To evaluate how well the reduced components preserved class-specific information, we applied the same decoding procedure to both the original and reconstructed data. For each time point, we compared the projection of each trial to the reconstructed class centroids (colour vs. scene) and assigned the label corresponding to the closest centroid. This produced a time-generalisation matrix analogous to that obtained from the original data.

We then computed the Spearman correlation between the decoding matrices derived from original and reconstructed data. This resulted in a two-dimensional matrix capturing, for each retrieval time point, how closely its reconstructed decoding profile matched the original profile across all encoding time points (See Supplementary Fig. 8c). The diagonal of this matrix reflects correspondence at matched time points. To summarise this relationship at the participant level, we averaged across retrieval time, yielding one correspondence value per participant and trial type.

These values showed a strong and significant correlation across participants (Spearman’s ρ = .87, p < .001), demonstrating that the PCA-based dimensionality reduction retained the majority of information relevant for distinguishing AM+ from AM- trials. Although decoding accuracy was used here as the validation metric, the same reconstruction procedure could be performed with any other measure of representational structure.

#### De-mixed Principal component analysis

The same procedures described previously were used for epoching, preprocessing, and ripple detection. We then divided the ripple-aligned data into the four target association conditions (red, blue, indoor, outdoor) and memory outcome (AM+ and AM-). For each participant, all extra-hippocampal channels were included. These data were then combined, resulting in a four-dimensional array (channels × target association × memory × time).

Data organisation followed the procedure outlined in ^61^ and made use of open-source code provided in ^96^. For analysis, default algorithm parameters were used. The marginalisation variables of interest were: (1) target association, (2) memory, (3) their interaction, and (4) an independent component. The lambda parameter was optimised by running 100 iterations using *dpca_optimizeLamda*, which yielded the decoder (w) and encoder (v) matrices for each component. In addition, the noise covariance matrix was estimated using *dpca_getNoiseCovariance*, and explained variance was quantified *dpca_explainedVariance*. This explained variance estimate, based on the noise covariance, enabled us to distinguish structured neural signals from random noise. Since all estimated signal was captured by the first 50 demixed principal components (dPCs), any remaining variance was likely attributable to noise (see Supplementary Fig. 11a) ^61^.

For decoding, we included the top 3 components for each marginalisation and repeated the classification 100 times using *dpca_classificationAccuracy*. To quantify how much task-relevant information was captured by the demixed components, we used the *dpca_classificationAccuracy* function from the original dPCA toolbox. This procedure performs cross-validated linear decoding on the dPCA projections. The function first computes the noise covariance matrix across conditions, then runs dPCA on the training data to obtain decoder (W) and encoder (V) matrices for each marginalisation (e.g. target association, memory, their interaction, and time). For each cross-validation repetition, dpca_classificationAccuracy randomly assigns a subset of trials to a test set and uses the remaining trials to form a training set (“pseudo-ensembles”). The training data are projected onto the selected dPCs, and for each marginalisation and component, class means are computed along the relevant task dimensions (here corresponding to the four target associations, the two memory outcomes, or their interaction). The test data are projected using the same decoders, and at each time point the function assigns each test observation to the nearest class mean in this one-dimensional projection (minimum absolute distance). Classification accuracy is then computed as the proportion of correctly assigned conditions, averaged across repetitions and normalised by the number of conditions.

The output is an accuracy matrix with dimensions (marginalisation × component × time), where each element reflects how well that particular dPC discriminates between the predefined classes at a given time point. In our analyses, we focused on the first three components for each marginalisation (numComps = 3) and used the default cross-validation settings (100 repetitions). To assess statistical significance, these empirical accuracy time courses were compared against a label-shuffled baseline using the same decoding pipeline (see below for details on the permutation-based cluster correction).

To obtain a baseline, label-shuffled data were used, with 500 shuffles repeated 100 times using *dpca_classificationShuffled*. Significant time windows were determined via 500 iterations of stratified Monte Carlo leave-group-out cross-validation. Trial labels were shuffled 500 times using a stratified approach to maintain an equal number of trials per condition. See ^61^ for full implementation details. The number of repetitions and shuffles was selected to provide stable estimates of the decoding distribution.

To assess statistically significant differences from the shuffled baseline, we applied a cluster-based permutation test using z-scored decoding accuracy values (note: this procedure is not part of the original dPCA toolbox but was added to control for multiple comparisons). Z-scores were computed by subtracting the mean of the shuffled distribution from the empirical decoding values and dividing by the standard deviation. Consecutive time points exceeding a z-threshold of ±1.96 (two-sided α = .05) were grouped into clusters using binary connected component labelling (*bwlabel* in MATLAB). For each cluster, the sum of z-values was computed as the test statistic. A null distribution was generated via 1000 random permutations of the accuracy time series, and the maximum cluster-level statistic was recorded for each permutation. Observed cluster statistics were compared to the 97.5th percentile of the null distribution, and clusters exceeding this threshold were deemed statistically significant. This procedure was conducted separately for each dPC. Because each marginalisation involved a different number of conditions (e.g., interaction: 8 classes, chance = 12.5%; memory: 2 classes, chance = 50%), decoding results in Fig. 3d are plotted as the empirical accuracy minus the shuffled baseline. In Supplementary Fig. 11c-g, both empirical and baseline decoding values are shown.

To visualise the components, we projected the original data onto the weight matrix and plotted the first dPCs corresponding to the target association × memory interaction, target association, and memory effects (dPCs 1, 2, and 4 in main Figure 3, and 1, 3, and 4 in Supplementary Fig. 11h). To visualise neural trajectories in state space, we multiplied each component by the original data and separated trials into pre- and post-ripple windows. For post-ripple, we included data from 400 to 800 ms after ripple onset, corresponding to the peak of dimensionality expansion. To ensure symmetry in comparison, we used −800 to −400 ms for the pre-ripple window.

The silhouette score, which measures how well-separated data points are within their assigned variables with values ranging from zero to one, where higher scores indicate stronger clustering, was calculated using MATLAB’s *silhouette* function. Statistical results for dPCs [1, 2, and 4], and [1, 3 and 4] are reported in the main text.

K-means clustering was performed for cluster counts ranging from 1 to 10 using MATLAB’s built-in *kmeans* function, with each k run 10 times using random initial centroids. K-means clustering identifies clusters by minimising the distance between data points and their respective cluster centres. We plotted the mean silhouette value across cluster sizes. The optimal cluster number was identified for both pre- and post-ripple windows. Statistical results for dPCs [1, 2, and 4] as well as [1, 3, and 4] are reported in the main text.

### Theta-gamma phase-amplitude coupling

The same procedure as for decoding and dimensionality estimation was used for epoching, processing and detecting ripples in the data, except that we now used a much broader time window of −4 to 6 seconds around ripple peak to account for later time-frequency transformation. We defined the peak frequency in the theta and gamma range, separately. To isolate oscillatory contributions ^47^ and to find the peak frequency of low (1-30Hz, in steps of 1Hz) and high (30-150Hz, in steps of 5Hz) frequencies, 1/f activity was attenuated in the time-frequency domain using the FOOOF algorithm ^100^ as implemented in the Fieldtrip toolbox ^101^. We then defined the peak theta frequency as the frequency with the highest power between 3 and 8Hz in hippocampal channels and for gamma frequency between 40 and 140Hz for all extra-hippocampal channels (to allow for ±2 and 10Hz for theta and gamma, respectively; see next). This procedure was done to ensure that the phase-amplitude coupling was performed on a narrow-band oscillation rather than broadband. Once defined we centred the data on these frequencies with a span for theta being ±2Hz and for gamma frequency being ±10Hz, with the peak frequency in the middle (frequency 0). Once having defined the peak frequency, the original data was subjected to another decomposition. Again, we divided high and low frequencies, using different methods to estimate phase and power. For low frequencies we convolved the data using a wavelet transformation with a hanning taper, with the number of cycles being roughly 500ms for each frequency, but never less than 5 cycles. For high frequencies, we estimated power using a multitaper method based on Slepian sequences as tapers. Frequency smoothing was set to one quarter of the frequency of interest and temporal smoothing was set to 200ms ^47^. The data were then baseline corrected between −500 to −100 pre-cue onset. The phase-amplitude coupling was performed per channel between −1 and 1 second, to comply with all other analyses. We performed the phase-amplitude coupling as in ^48,102^. For each phase-amplitude sample, we also ran a permuted baseline, where we shuffled the trials 500 times. The resulting data show the contrast between empirical and shuffled phase-amplitude coupling with the peak frequency for theta and gamma as 0 on the x and y-axis, respectively. The contrast was statistically tested by running a two-sided cluster-based permutation. For clarity, we note that our PAC computation reflects the phase-amplitude vector length (theta phase × gamma-band amplitude) relative to a trial-shuffled baseline, and the goal of this analysis is not to distinguish between gamma and ripple sub-bands per se, but to establish a temporal link between hippocampal ripples, hippocampal theta, fast cortical activity, and the subsequent expansion of cortical representational dimensionality.

To assess whether using region-level peak frequencies (hippocampal theta, cortical gamma) was justified, we quantified the dispersion of per-channel peak frequencies around each participant’s region-level peak. Hippocampal theta frequencies showed very low variability (mean SD = 1.88 Hz; MAD = 0.17 Hz), with 74.8% of hippocampal channels falling within ±2 Hz of the participant-level peak, indicating that hippocampal theta oscillations were strongly clustered around a narrowband frequency within each participant.

Cortical gamma peaks exhibited broader dispersion, as expected for gamma-band activity across distributed cortical regions (mean SD = 15.9 Hz; MAD = 0.42 Hz). The small MAD indicates that most channels nevertheless lie close to the participant-level gamma peak, with the larger SD driven by a minority of high-frequency outliers. Even so, 31.1% of channels fell within ±10 Hz of the regional peak, consistent with substantial regional variability in cortical gamma reported in prior work (e.g.,^66^). These findings justify our use of region-level theta and gamma peaks for PAC alignment while also validating the range of frequencies used in subsequent analyses.

Because non-sinusoidal high-frequency waveforms can artificially inflate phase-amplitude coupling, particularly if sharp transients differ across cortical regions, we performed additional control analyses quantifying gamma waveform shape for each cortical channel at its PAC-defined peak frequency. Following established approaches, we band-pass-filtered each channel around its participant-specific gamma peak (±10 Hz) for ripple-aligned trials, and extracted several morphological metrics. (1) peak-trough sharpness ratio, (2) rise-decay asymmetry, (3) waveform skewness, and (4) kurtosis.

For each cortical channel, we designed a 4th-order zero-phase Butterworth band-pass filter centred on the channel’s peak gamma frequency (±10 Hz), constrained to 30-150 Hz. Ripple-locked trials were then filtered and concatenated in time to yield a single continuous gamma-band time series per channel. On this concatenated signal, we detected local maxima (peaks) and minima (troughs). Peak and trough sharpness were computed by comparing each extremum to its local neighbourhood: for a given peak (or trough), we took the difference between its amplitude and the mean amplitude of k neighbouring samples on either side (here k = 3 samples), and then averaged across all peaks or troughs within the channel. A peak-trough sharpness ratio was defined as the mean peak sharpness divided by the mean trough sharpness.

To quantify rise-decay asymmetry, we measured the number of samples from each trough to the subsequent peak (rise time) and from each peak to the subsequent trough (decay time), and then took the ratio of mean rise to mean decay time for that channel. Thus, values >1 indicate longer rises than decays, and values <1 the opposite.

Finally, we characterised the overall distribution of the gamma-band signal using its skewness and kurtosis (third and fourth central moments), computed on the concatenated filtered time series for each channel. Together, these metrics (peak-trough sharpness ratio, rise-decay ratio, skewness, kurtosis) were used to quantify gamma waveform symmetry and asymmetry across cortical sites and to test whether systematic waveform differences could account for the observed theta-gamma PAC effects.

Gamma waveforms were highly symmetric across channels (peak-trough ratio: 1.000 ± .0045, mean ± SD; rise-decay ratio: 1.000 ± .0021), exhibited negligible skewness, and showed kurtosis values consistent with narrowband filtering (5.98 ± 14.04). Importantly, none of these metrics correlated with the preferred gamma frequency across channels (all p > 0.17), indicating that regional variation in gamma frequency or waveform shape cannot account for the observed PAC. Because the trial-shuffled surrogate preserves gamma waveform morphology while disrupting cross-trial hippocampal-cortical alignment, these results jointly confirm that our TG-PAC effects reflect genuine inter-areal coupling rather than artefacts of waveform asymmetry (Supplementary Fig. 12).

To assess when in time around the ripple events the TG-PAC was strongest, we ran the same analysis, but now binning the data into 500ms bins, with 90% overlap and only including peak gamma ±5Hz. We selected the length of the time window to allow for a minimum of one full cycle for each frequency before and after the ripple event (−1 to 1), constraining the binning to at least 500ms (1Hz frequency needs 1 second for a full cycle). Again, we ran a two-sided cluster-based permutation to test for significance between empirical data and a shuffled baseline (now between -.5 to 1 sec as we were mainly interested in the post-ripple effect). In a subsequent analysis, we correlated the dimensionality transformation with the TG-PAC using a two-sided Spearman’s correlation.

Lastly, to understand the temporal directionality, we performed a cross-correlation between TG-PAC and dimensionality expansion. Due to difference number of sample points for the two vectors, we linearly interpolated them using MATLAB’s *interp1* function. For each participant, we then generated 1000 null-distributions. To keep the temporal autocorrelation in the surrogate data, we used the Iterative Amplitude Adjusted Fourier Transforms ^103^, which instead of randomly shuffling time points, shuffles phase-values. We then z-scored the observed group-average cross-correlation against the distribution of permuted cross-correlations at each time lag. To correct for multiple comparisons across lags, we applied a cluster-based approach: z-score values exceeding a threshold of 1.96 (alpha .05 for two-sided test) were binarised, and temporally contiguous clusters of supra-threshold points were identified using connected component labeling (*bwlabel* in MATLAB). The sum of z-scores within each cluster was computed as a cluster-level statistic, and the maximum cluster sum across the entire lag window was retained for the real data. This procedure was repeated for each of the 500 permutations to generate a null distribution of maximum cluster statistics. A p-value was computed by comparing the real maximum cluster statistic to this null distribution, quantifying the probability that a cluster of equal or greater strength would be observed under the null hypothesis.

### Conducting analyses based on spikes instead of ripples

We conducted the reinstatement and dimensionality analyses aligned to interictal epileptiform discharges (IEDs) rather than ripples to rule out that our results could be driven by epileptiform artifacts. To identify IEDs, we applied a previously used outlier-based spike detection approach (Honey et al., 2011), in which transient high-amplitude deflections are detected as samples exceeding a robust amplitude threshold relative to each channel’s distribution. Specifically, for each hippocampal channel, we computed the channel median and interquartile range (IQR) over the analysed recording segment and flagged candidate spike samples whose absolute deviation from the median exceeded 2 × IQR, considering both positive and negative deflections. To remove the detected events from the continuous signal, candidate spike samples were temporally dilated by ±10 ms and the affected samples were replaced using piecewise cubic interpolation from the surrounding non-spike samples. For alignment analyses, we collapsed each suprathreshold segment to a single peak sample by selecting, within each contiguous suprathreshold run, the sample with maximal absolute deviation from the channel median. To avoid counting high-frequency burst activity as multiple spikes, we imposed a 100 ms within-channel refractory period, retaining only the first detected peak within each refractory window. Spike-triggered averages were computed by extracting ±50 ms epochs around each spike peak. Epochs were high-pass filtered at 5 Hz and baseline-corrected by subtracting the mean voltage in a pre-spike window (−100 to −20 ms) prior to averaging across spikes and channels.

First, we computed spike-triggered averages from hippocampal channels (Supplementary Fig. 14). We found numerically more spikes in AM- trials (M = 174, SE = 69) as compared to AM+ trials (M = 117, SE = 30), but no difference between number of spikes (t(11) = −1.08, p = .30. This indicates that epileptiform activity is not statistically more elevated on AM+ vs. AM- trials. We then repeated the full reinstatement analysis (Fig. 3a), aligning data to spike peaks instead of ripple peaks. No significant cluster emerged for spike-aligned reinstatement, contrasting AM+ and AM- trials (Supplementary Fig. 15a). This stands in sharp contrast to the robust reinstatement observed when aligning to ripples. We also repeated the PCA-based dimensionality analyses (Fig. 3b) using spike alignment. Again, we found no significant increase for AM+ trials compared to AM- trials (Supplementary Fig. 15b). Finally, we repeated the analyses presented in Fig. 3a-b while excluding ripples that occurred within ±500 ms of a detected spike in any hippocampal channel (Supplementary Fig. 6a-b). The results remained qualitatively similar.

Across all analyses, number of spikes for AM+ and AM-, reinstatement and PCA epileptic spikes failed to produce any of the neural signatures we observe around ripples. This demonstrates that the ripple-locked effects in reinstatement and dimensionality cannot be attributed to epileptiform artefacts.

---

**Supplementary Figure 1.**
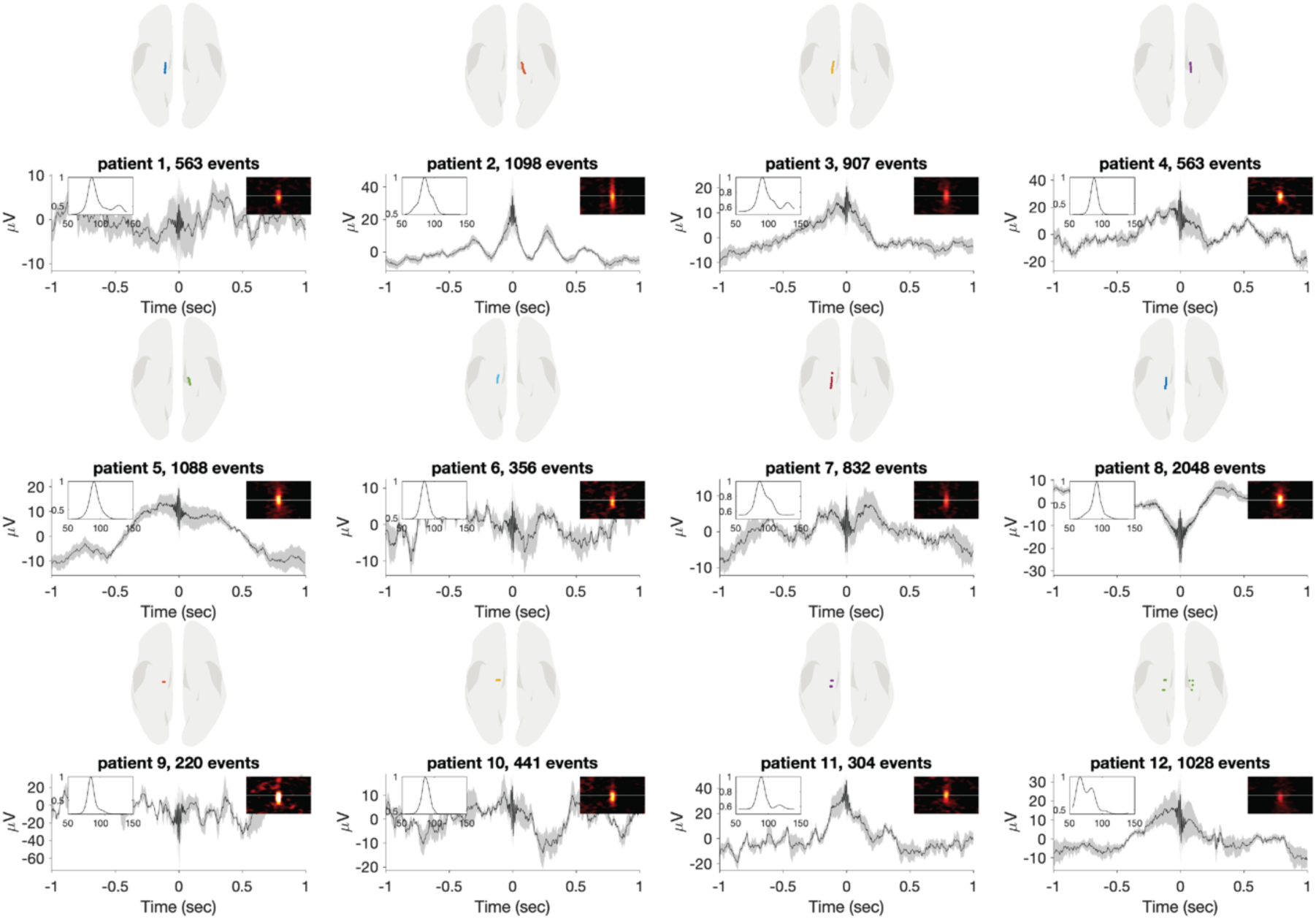
Electrode implantation, ripple counts, and spectral characteristics per participant. Each panel displays data from a single participant. The top row illustrates electrode implantation sites within the hippocampus. The middle row shows the number of detected ripple events. The bottom row presents the average ripple waveform alongside the corresponding time-frequency and spectral power profile.

**Supplementary Figure 2.**
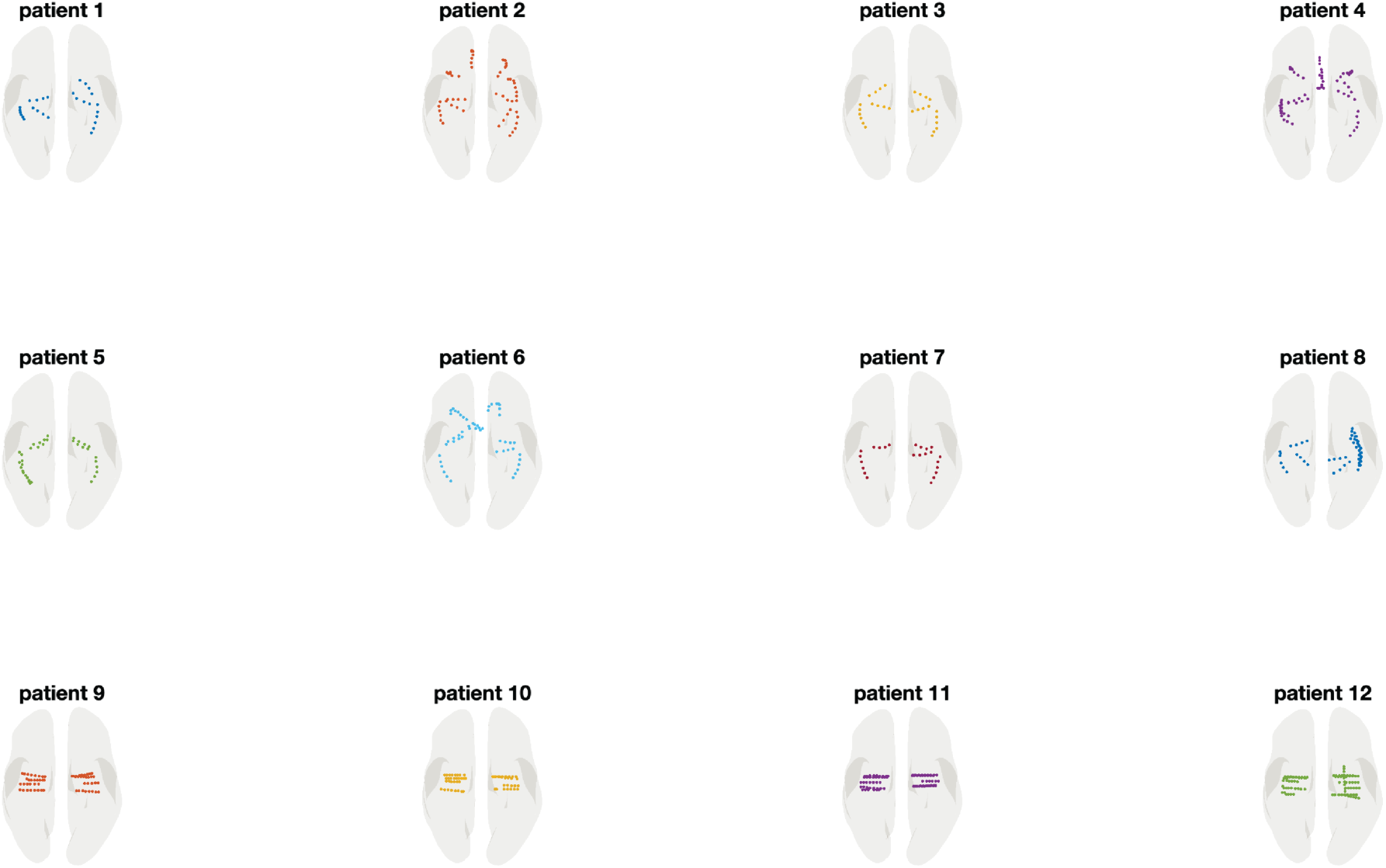
Electrode implantation extra-hippocampal regions.

**Supplementary Figure 3.**
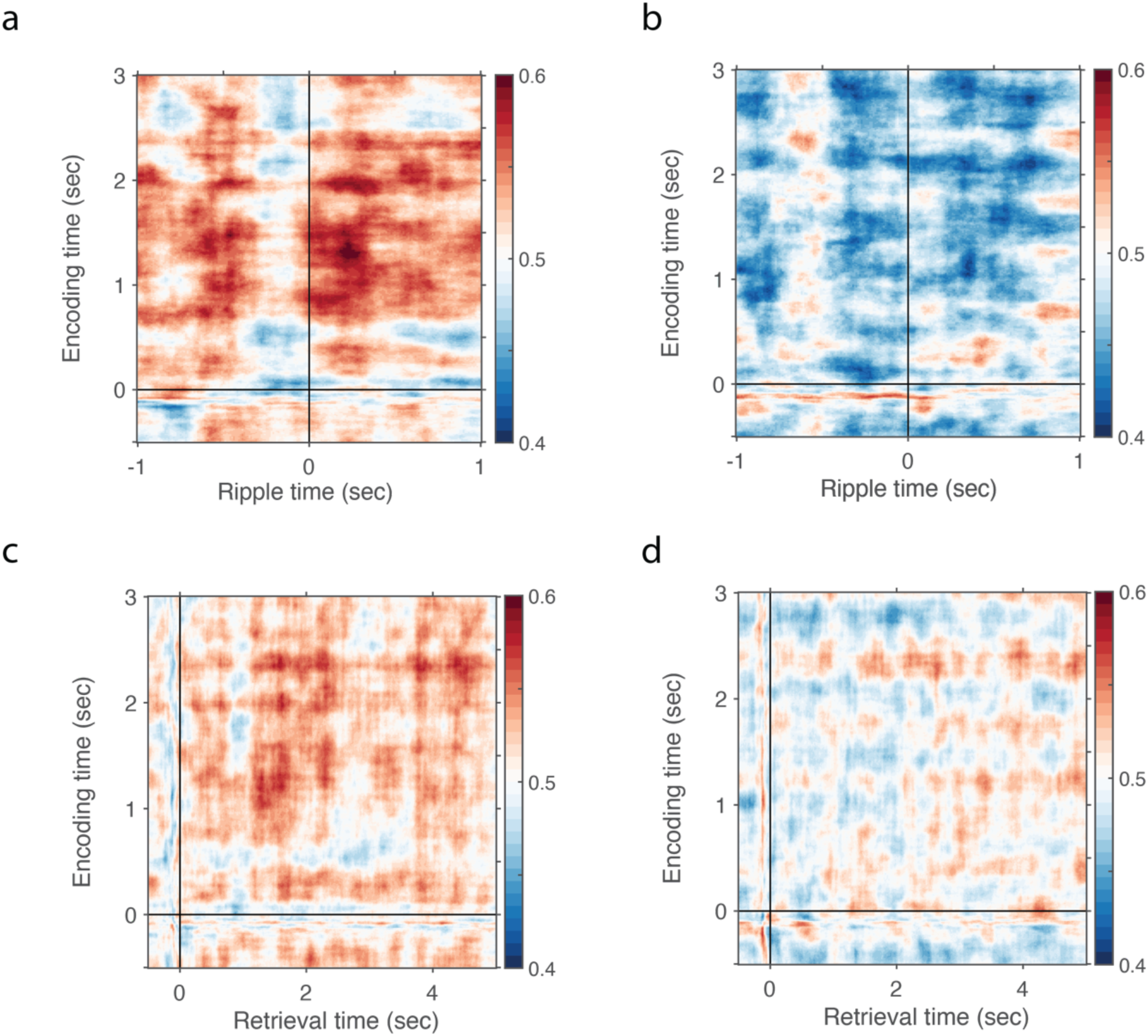
Reinstatement ripple-aligned and cue-aligned for AM+ and AM-, separately. **a** AM+ trials showed highest decoding accuracy following ripple events. **b** Chance-level decoding around ripple events for AM- trials. **c** When aligning the data to cue-onset at retrieval, we found the highest decoding accuracy around 1.5 to 2.5 seconds during encoding and retrieval. **d** Chance-level decoding during the entire retrieval period for AM- trials.

**Supplementary Figure 4.**
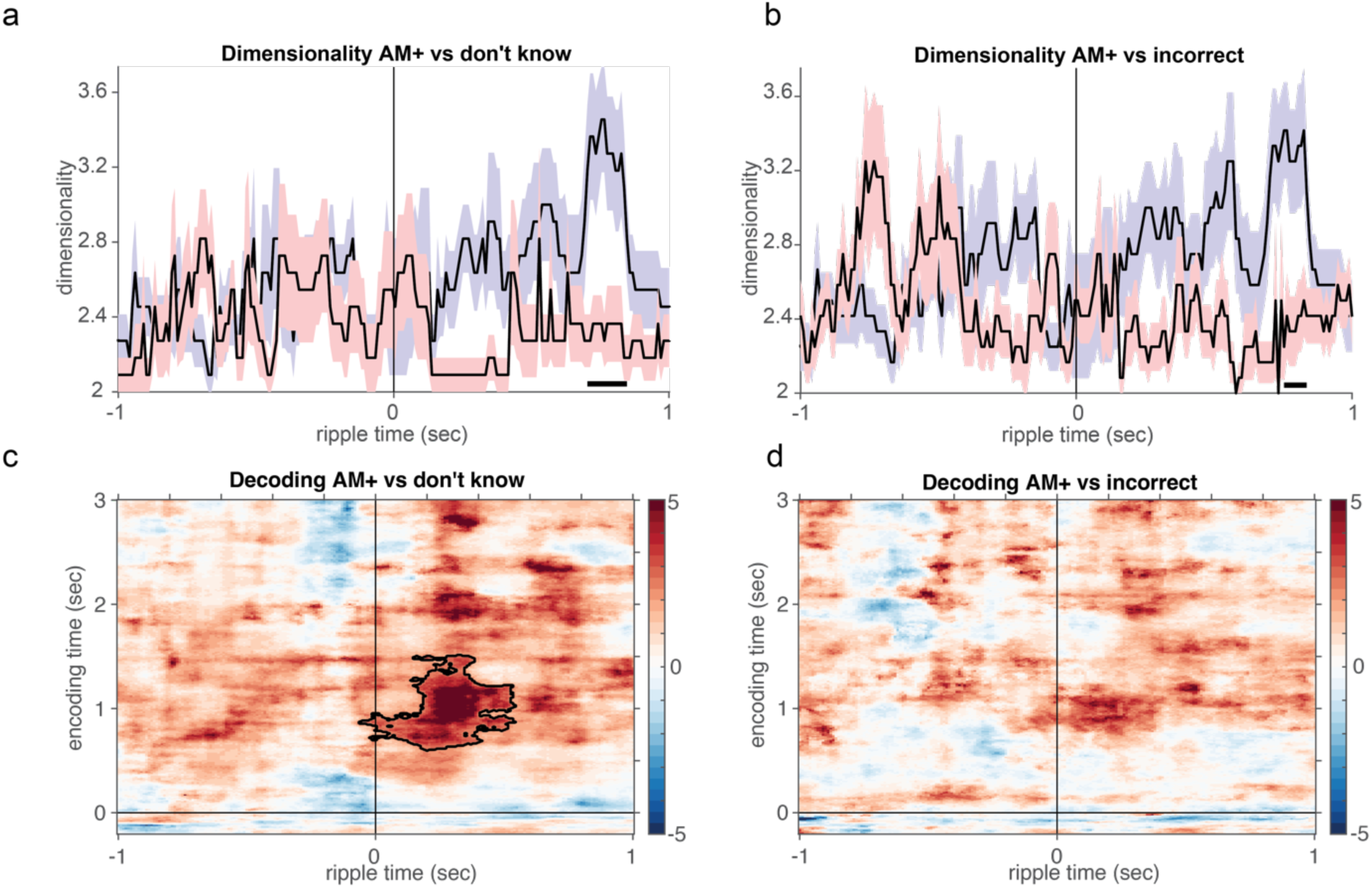
Dimensionality and reinstatement analyses dividing AM- into incorrect and don’t know responses. **a** AM+ trials showed significant higher dimensionality as compared to don’t know responses in a similar time window as main analysis in Fig. 3b. **b** AM+ trials showed significant higher dimensionality as compared to incorrect responses in a similar time window as main analysis in Fig. 3b. **c** AM+ trials showed significant higher decoding accuracy as compared to don’t know responses in a similar time window as main analysis in Fig. 3a. **d** AM+ trials showed numerically higher decoding accuracy as compared to incorrect responses in a similar time window as main analysis in Fig. 3a.

**Supplementary Figure 5.**
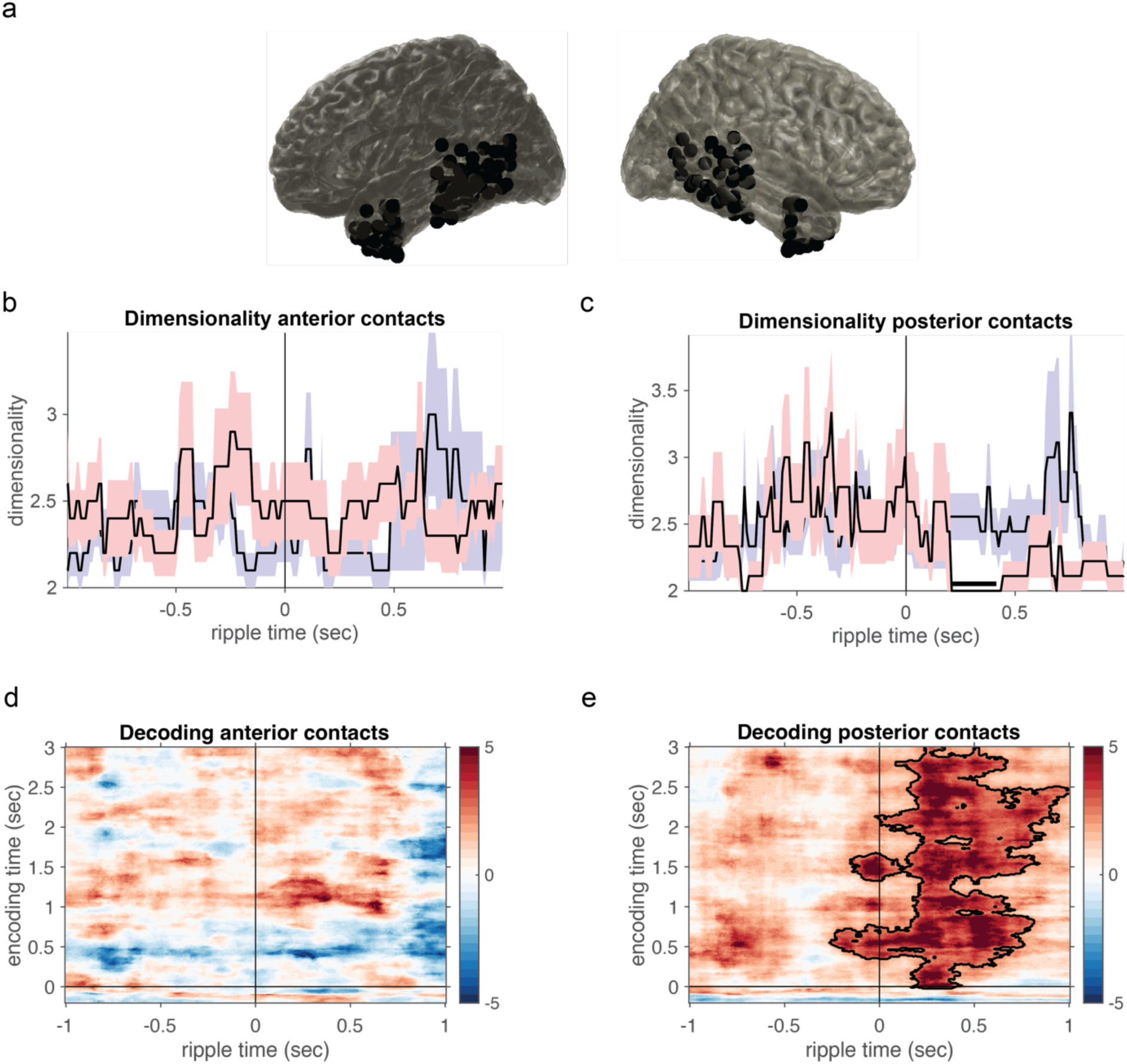
Dimensionality and decoding analyses using anterior and posterior cortical contacts. ***(A)*** Included anterior and posterior contacts. **(B)** No significant difference between AM+ and AM- for anterior contacts, but with the same increase around 600-800ms as in Fig 3a. **(C)** Significant difference between AM+ and AM- in a similar time window as in Fig. 3a. **(D)** No significant decoding difference between AM+ and AM- for anterior contacts, but with largest difference in the same post-ripple time window as in Fig. 3a. **(E)** Significant difference post ripple between AM+ and AM- for posterior contacts.

**Supplementary Figure 6.**
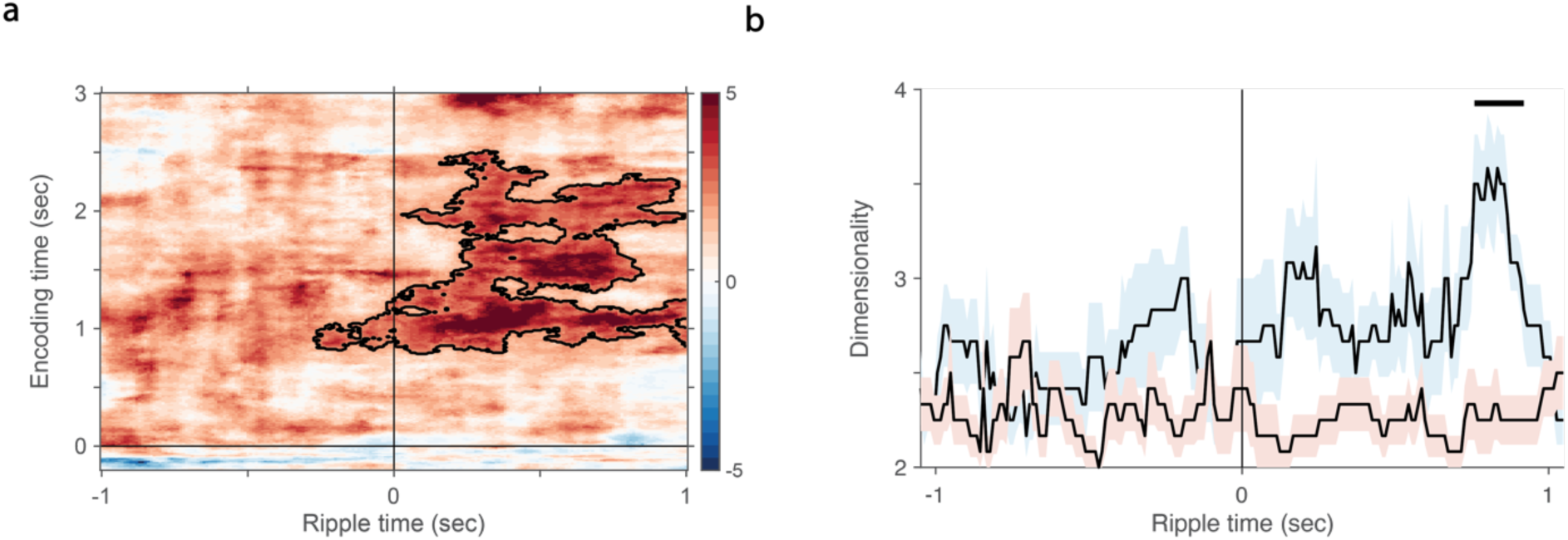
Spike-corrected reinstatement and dimensionality. **a** Reinstatement analysis excluding ripples occurring within ±500 ms of spikes in hippocampal channels revealed a similar significant cluster as in Fig. 3a. **b** The spike-corrected time-resolved dimensionality analysis showed a comparable significant cluster around 400–800 ms post-ripple, consistent with Fig. 3b.

**Supplementary Figure 7.**
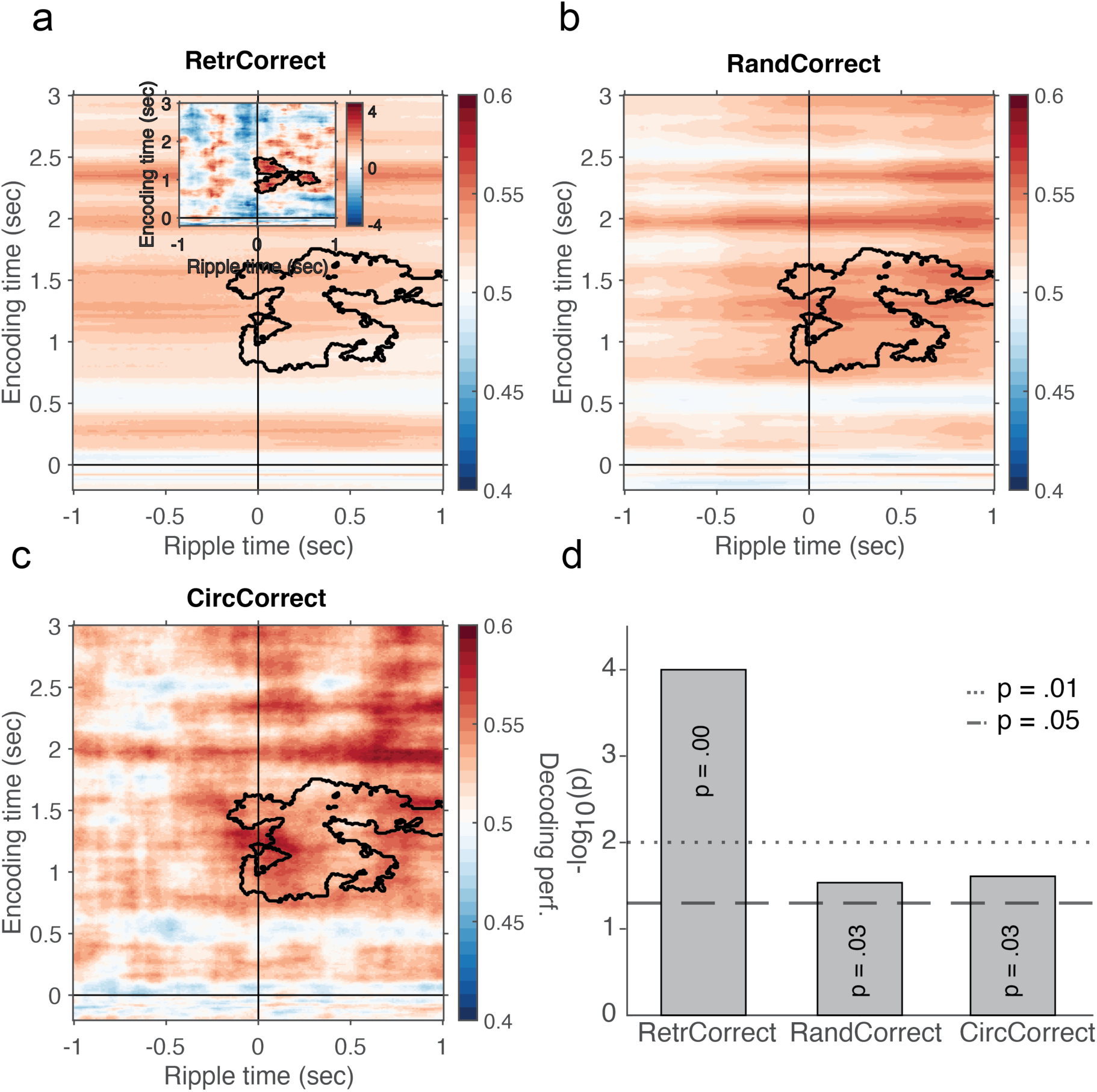
Control analyses confirming ripple-triggered reinstatement specificity. **(A)** From 1-5 seconds after cue onset, we picked 60 time points and treated them as ripple events and performed the same analysis as in 3a. The time generalisation matrix shows the decoding accuracy for these events with the significant cluster identified in Figure 3a highlighted in black. Inset shows a cluster-based permutation between the permuted baseline and empirical data, with a significant cluster in the same time window as in Figure 3a. **(B)** Decoding accuracy when AM+ trials were shuffled 1000 times. **(C)** A similar analysis was performed using a stricter control in which ripple times were swapped with those of adjacent trials (i.e., neighbouring trial shuffling). This revealed significant higher decoding in the empirical AM+ condition compared to the adjacent-trial control. **(D)** P-values when comparing empirical data with the different controls.

**Supplementary Figure 8.**
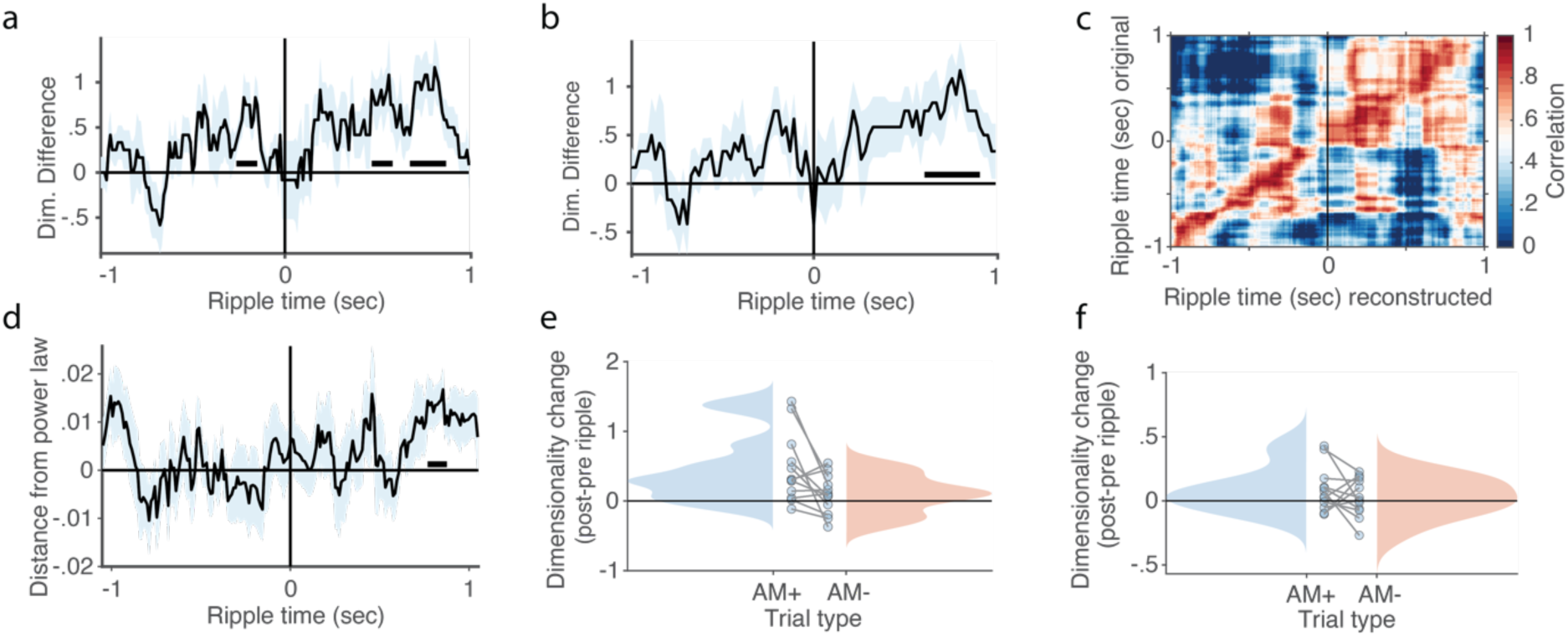
Robustness of dimensionality estimate, reconstruction analysis and alternative methods. **a** Using a 100 ms sliding window for dimensionality estimation produced results highly similar to those obtained with a 60 ms window, demonstrating the robustness of the approach. **b** same as **(a)** but with a 200 ms sliding window. **c** Correlation between original and reconstructed data, using the same number of components as in the PCA analysis, revealed a strong diagonal pattern, indicating a high-fidelity reconstruction of the original data structure. **d** Calculating the distance to a power-law distribution provided similar results as in 3b. **e** Estimating dimensionality using effective dimensionality provided similar results as in 3c. **f** Estimating dimensionality using Marčenko-Pastur threshold provided similar results as in 3c.

**Supplementary Figure 9.**
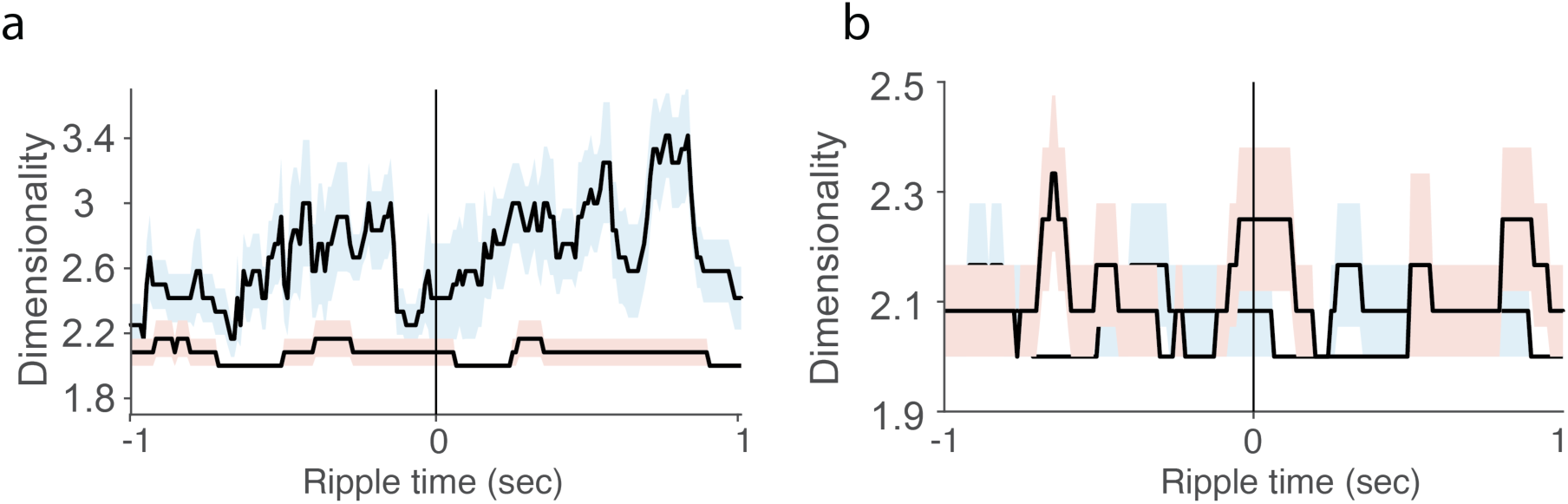
Dimensionality hippocampal vs. extra-hippocampal channels. **a** Extra-hippocampal channels (blue) had higher dimensionality as compared to hippocampal channels (red) for AM+ trials. **b** No difference between AM+ (blue) and AM-(red) trials in hippocampal channels. Note that, PCA on few channels in hippocampus will naturally not be very informative.

**Supplementary Figure 10.**
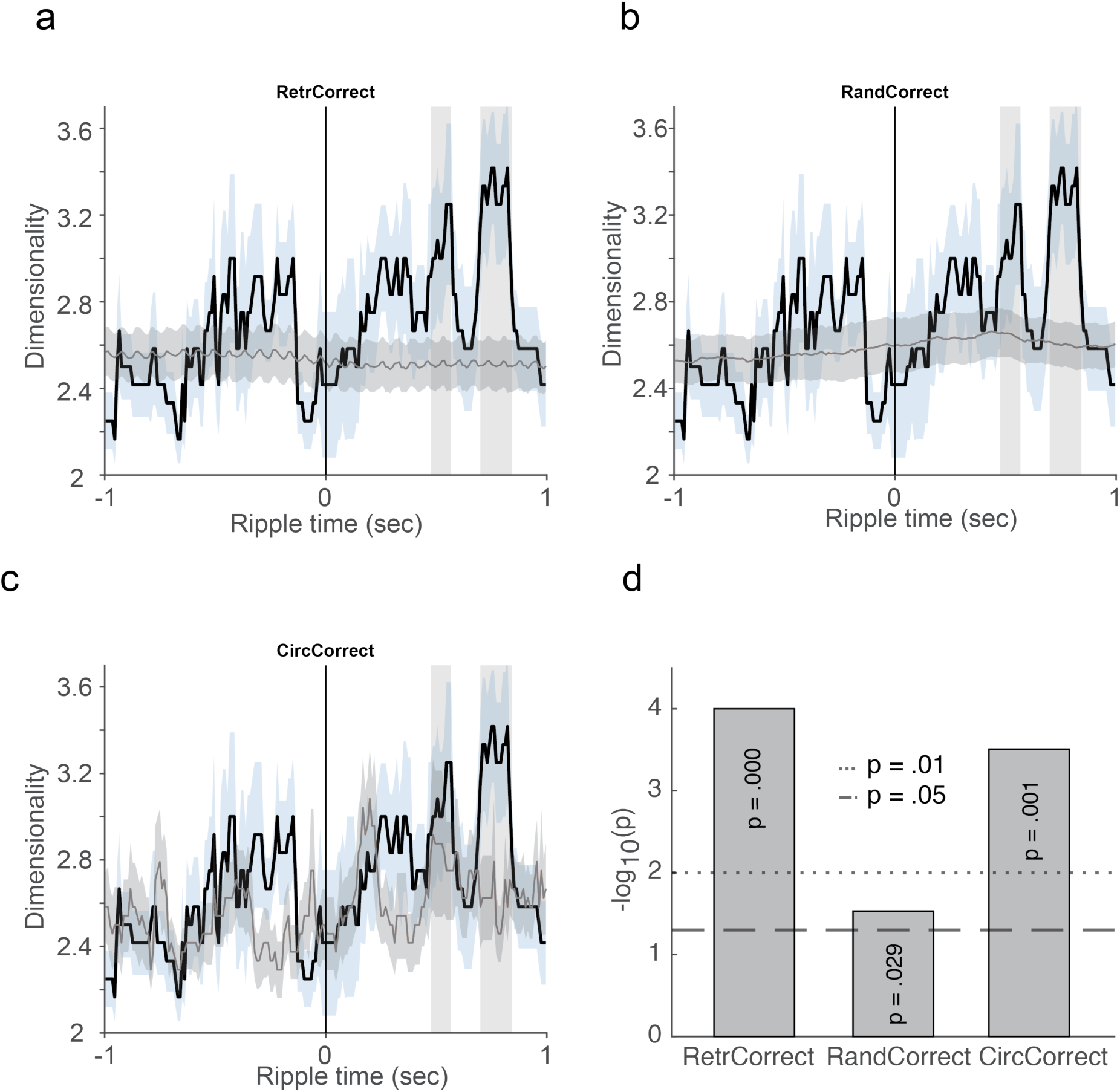
Control analyses confirming specificity of dimensionality expansion. **a** From 1-5 seconds after cue onset, we picked 60 time points for each participant and treated them as ripple events and performed the same analysis as in 3b. Grey is permuted baseline, blue is empirical data. **b** Shuffling AM+ trials 1000 times. Grey is permuted baseline; blue is empirical data. **c** Empirical AM+ (blue) and adjacent trials (i.e. temporally shifted controls, grey). **d** P-values when comparing empirical data with the different controls.

**Supplementary Figure 11.**
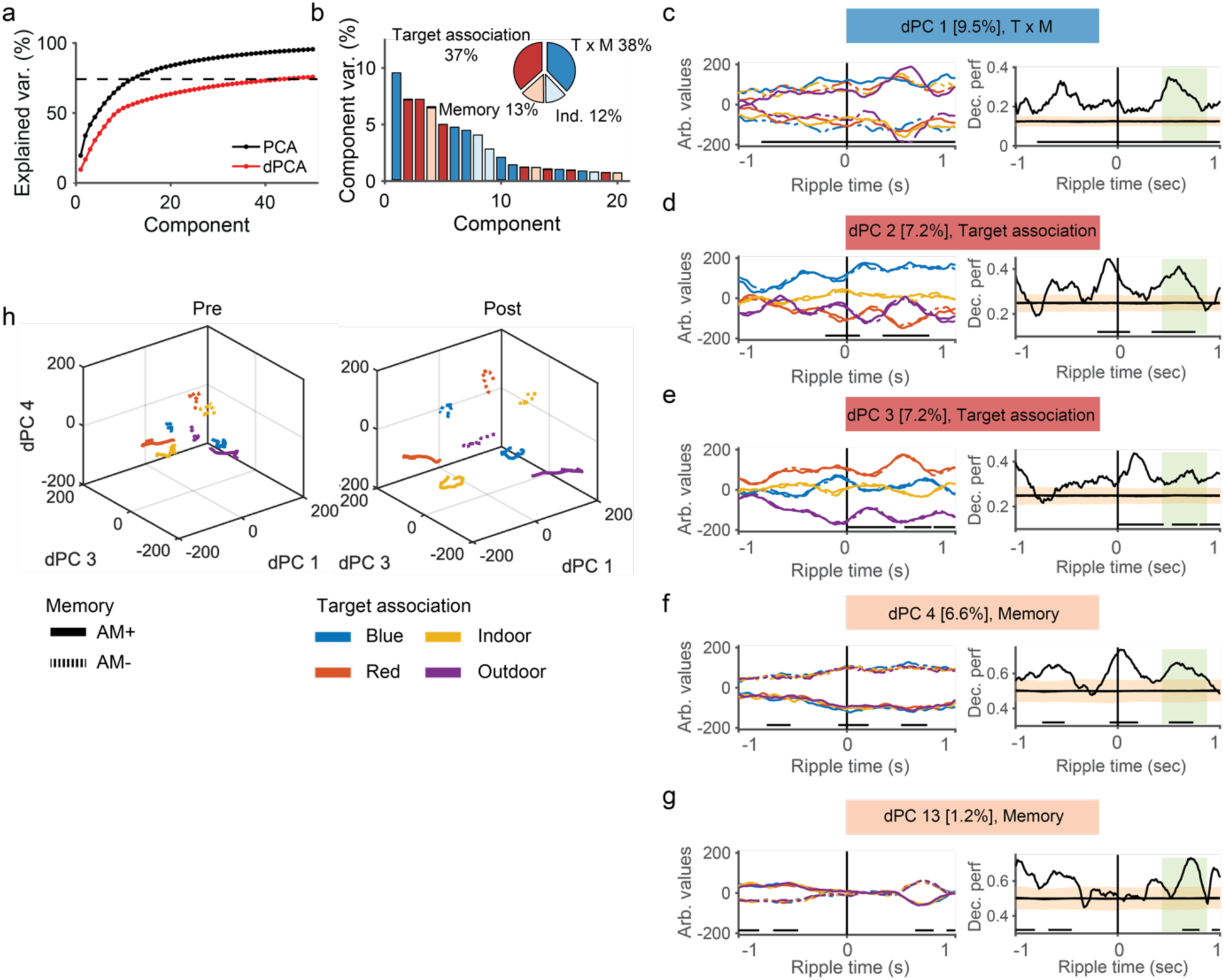
Demixed principal component analysis (dPCA). **a** Cumulative variance explained by the first 50 components exceeded the estimated signal variance (dashed line). Standard PCA (black) accounted for more total variance than dPCA. **b** The first four dPCs primarily reflected a mixture of “target association” and “memory” (dark blue), “target association only” (red), and “memory only” (beige). **c-g** Decoding accuracy for individual components, shown only for those with significant decoding between 400-800 ms post-ripple peak (highlighted by green bars). Left panels display component activation; right panels show decoding performance. Significant decoding intervals are marked with black horizontal lines. Orange shaded areas represent the shuffled distributions, with their mean indicated in black. **h** Neural state space plots of dPCs 1, 3, and 4 show that pre-ripple representations were more overlapping (left), whereas post-ripple activity exhibited greater separation between experimental conditions (right), indicating increased representational differentiation.

**Supplementary Figure 12.**
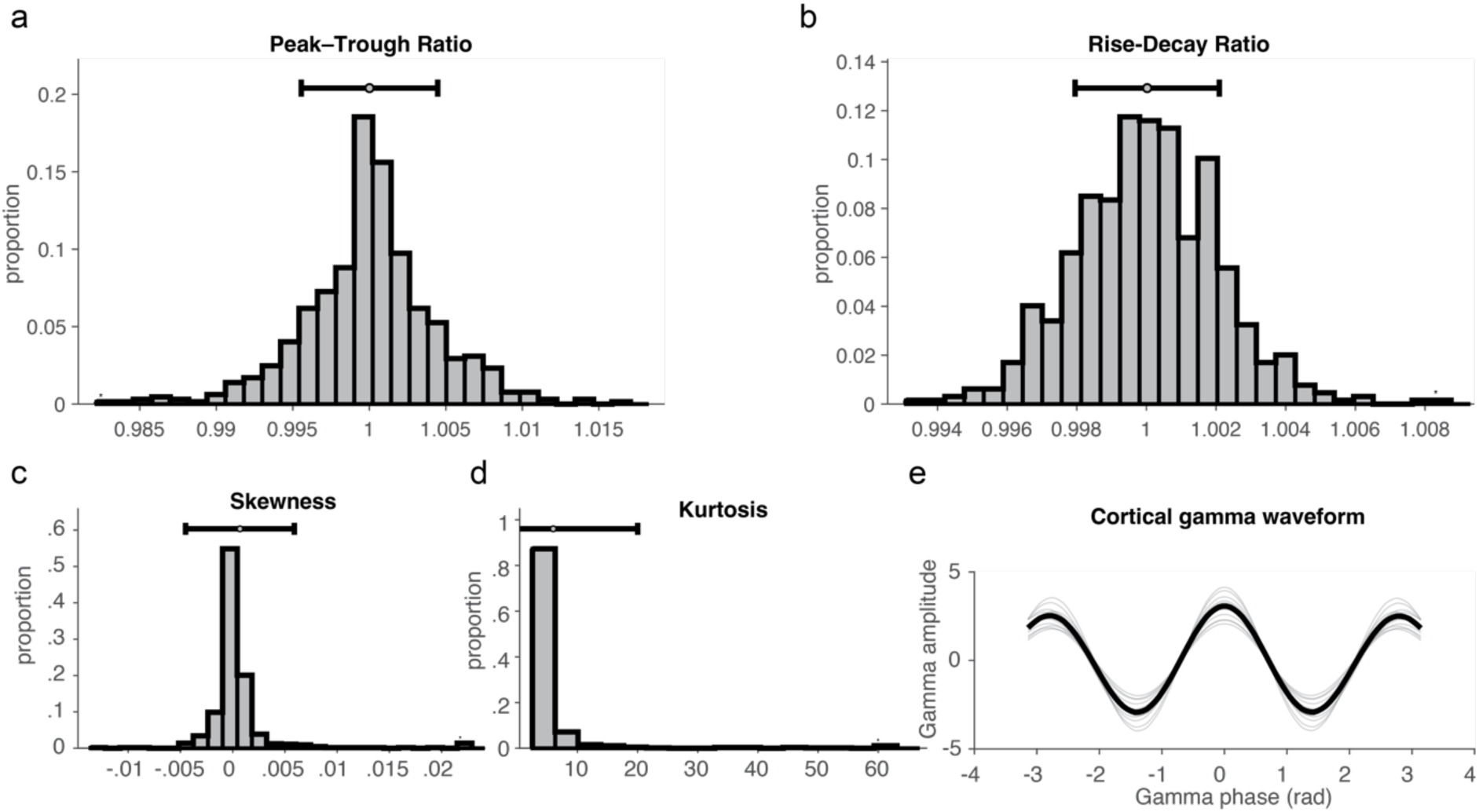
Gamma waveform-shape metrics. **a** Peak-trough sharpness ratio: ratio of mean peak sharpness to mean trough sharpness, reflecting symmetry of extrema. Values tightly clustered around 1 indicate near-perfect symmetry. **b** Rise-decay ratio: ratio of mean rise time (trough to peak) to mean decay time (peak to trough). Ratios near 1 indicate minimally asymmetric cycles. **c** Skewness of the gamma-band waveform, indexing asymmetry of the signal’s amplitude distribution (values centred at zero indicate symmetric waveforms). **d** Kurtosis, reflecting the heaviness of the tails of the amplitude distribution. The distribution is consistent with narrowband-filtered gamma activity, and no systematic deviations were observed. **e** The average cortical gamma waveform.

**Supplementary Figure 13.**
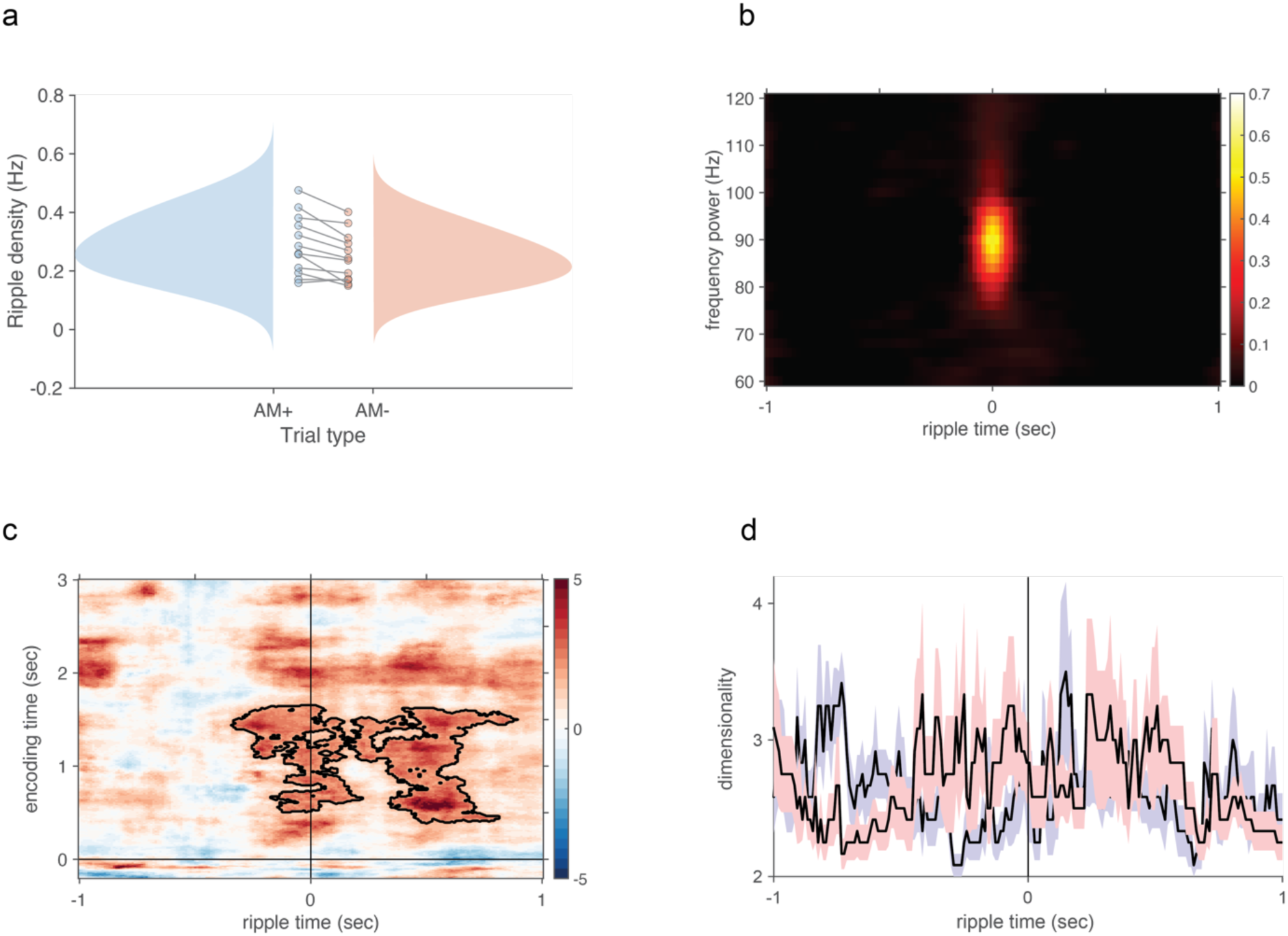
Ripple density, Spectral peak of ripples, Reinstatement and dimensionality for bipolar referencing. ***a*** Ripple density was higher for AM+ vs. AM- trials **b** Peak of ripples at 90Hz. **c** Time-resolved dimensionality analysis did not show the same significant cluster around 400-800ms post ripple, which is to be expected under bipolar reference scheme.

**Supplementary Figure 14.**
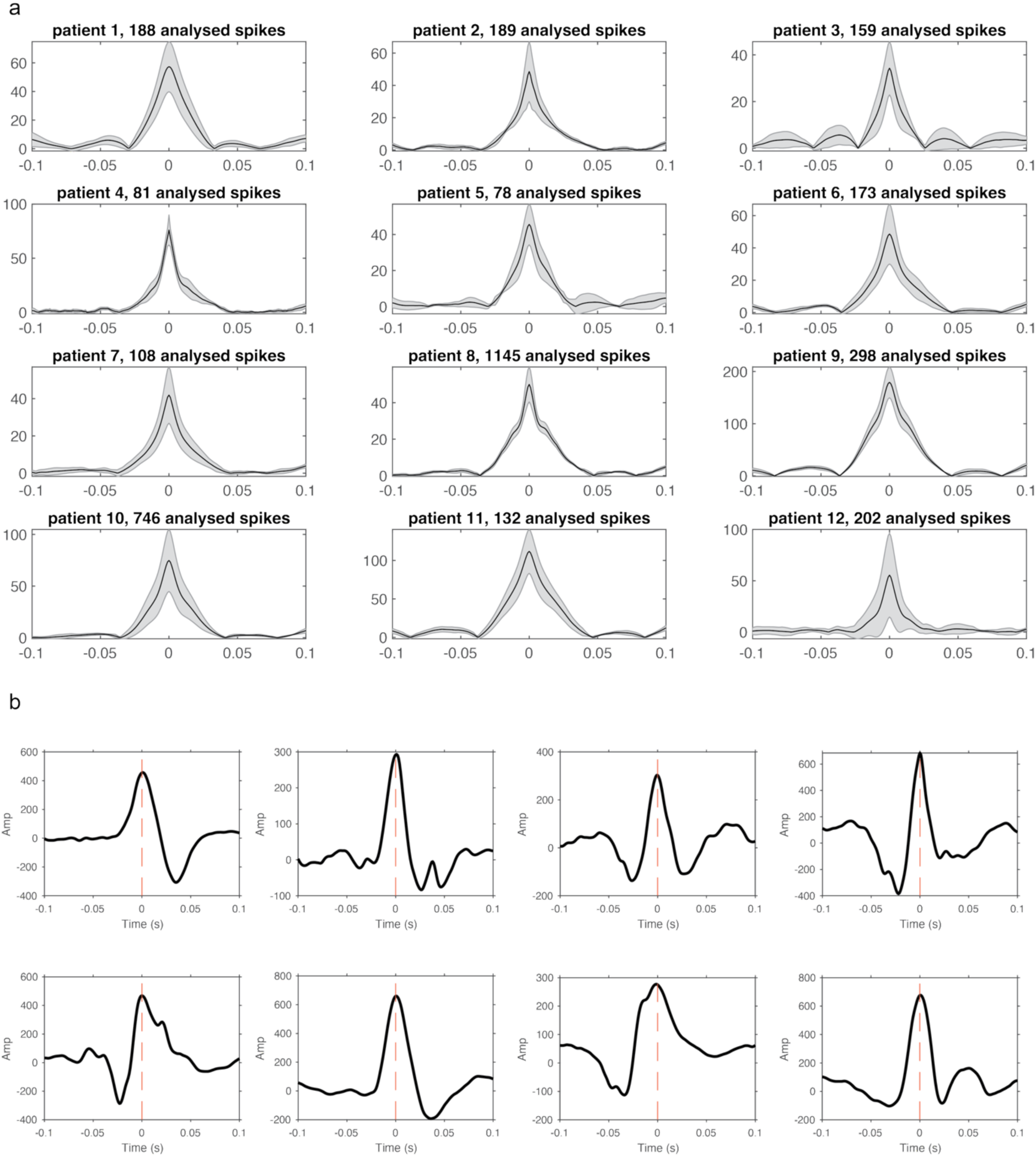
Spike-triggered averages. ***a***. For each participant, we extracted and averaged spikes, with the number of spikes above each spike-triggered average. **b.** Example of extracted spikes.

**Supplementary Figure 15.**
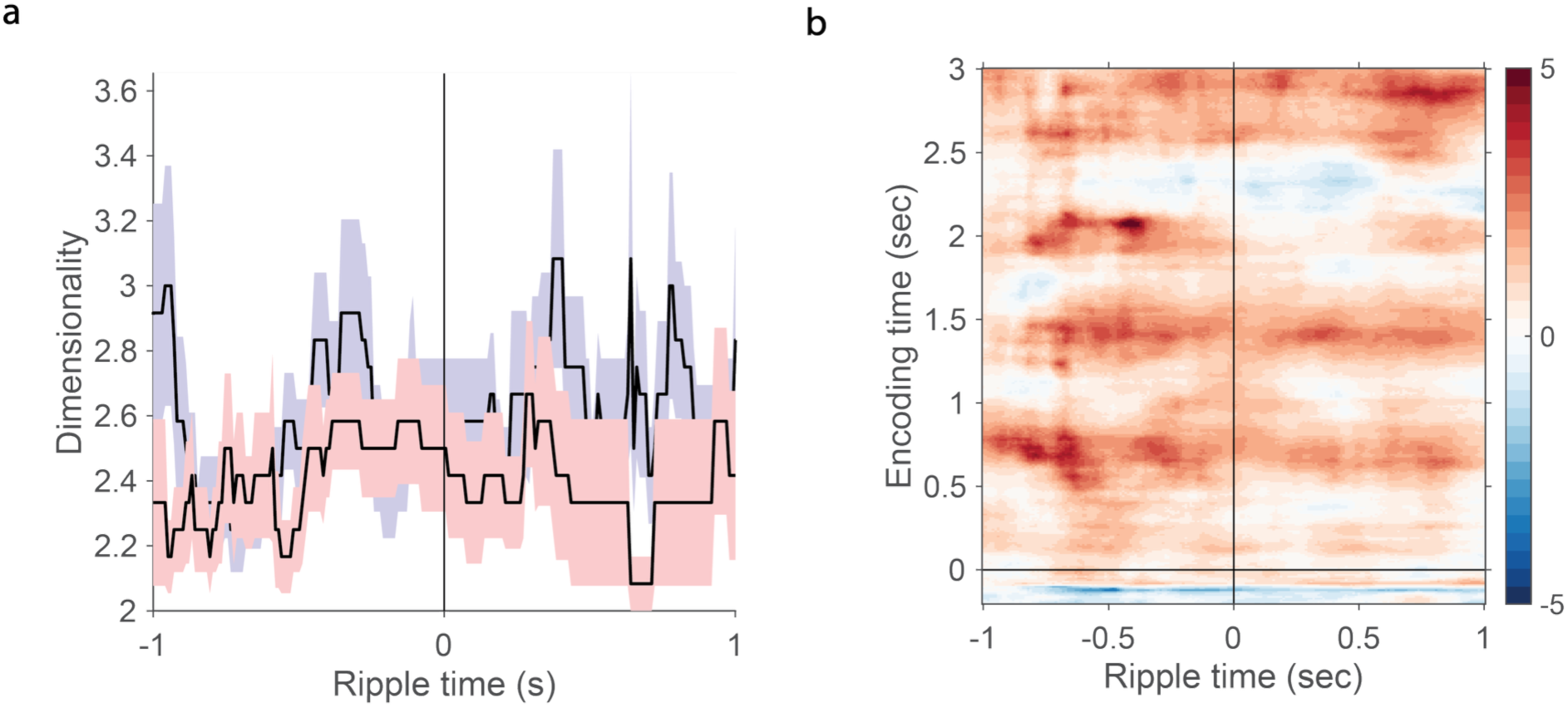
Dimensionality and decoding analyses based aligned on spikes. **a** Dimensionality analysis as in Fig. 3b showed no difference between AM+ (blue) and AM- trials (red). **b** Decoding analysis as in Fig. 3b showed no difference between AM+ and AM- trials.

**Supplementary Table 1.** Characteristics per patient. Number of analysed trials is for one ripple per trial.

| Patient | Implantation scheme | Number of analysed HC contacts | Number of analysed extra-HC contacts | Number of ripples | Number of analysed AM+ trials | Number of analysed AM- trials |
| --- | --- | --- | --- | --- | --- | --- |
| 1 | Longitudinal | 8 | 28 | 563 | 51 | 63 |
| 2 | Longitudinal | 7 | 59 | 1098 | 34 | 89 |
| 3 | Longitudinal | 7 | 28 | 907 | 48 | 109 |
| 4 | Longitudinal | 6 | 70 | 563 | 81 | 27 |
| 5 | Longitudinal | 6 | 37 | 1088 | 144 | 45 |
| 6 | Longitudinal | 6 | 53 | 356 | 48 | 46 |
| 7 | Longitudinal | 8 | 24 | 832 | 35 | 41 |
| 8 | Longitudinal | 7 | 54 | 2048 | 44 | 115 |
| 9 | Lateral | 2 | 61 | 220 | 31 | 29 |
| 10 | Lateral | 3 | 59 | 441 | 59 | 63 |
| 11 | Lateral | 4 | 81 | 304 | 42 | 19 |
| 12 | Lateral | 10 | 93 | 1028 | 87 | 38 |

**Supplementary Table 2.** Linear mixed-effects model of dimensionality transformation.

| Name | Estimate | SE | t | df | p | 95% CI (lower) | 95% CI (upper) |
| --- | --- | --- | --- | --- | --- | --- | --- |
| Intercept | 2.28 | .07 | 32.47 | 19,988 | <.001 | 2.14 | 2.42 |
| Trial Type | .09 | .03 | 2.82 | 19,988 | .005 | .03 | .16 |
| Half | .14 | .03 | 4.39 | 19,988 | <.001 | .08 | .21 |
| Trial Type x Half | -.09 | .02 | -4.16 | 19,988 | <.001 | -.13 | -.05 |

## Notes

### Competing Interest Statement

The authors have declared no competing interest.

### Summary of Updates

Substantial revision based on reviewers' comments. Additional supplemental figures etc.

## References

1. Tulving, E. Elements of Episodic Memory. (Oxford University Press, 1983).

2. Almeida, L. de, Idiart, M. & Lisman, J. E. Memory retrieval time and memory capacity of the CA3 network: Role of gamma frequency oscillations. Learn. Mem. 14, 795–806 (2007).

3. Bates, C. J. & Jacobs, R. A. Efficient data compression in perception and perceptual memory. Psychological Review 127, 891–917 (2020).

4. Barlow, H. B. Possible Principles Underlying the Transformations of Sensory Messages. in Sensory Communication (ed. Rosenblith, W. A.) 216–234 (The MIT Press, 2012). doi:10.7551/mitpress/9780262518420.003.0013.

5. Kerrén, C., Reznik, D., Doeller, C. F. & Griffiths, B. J. Exploring the role of dimensionality transformation in episodic memory. Trends in Cognitive Sciences https://doi.org/10.1016/j.tics.2025.01.007 (2025) doi:10.1016/j.tics.2025.01.007.

6. Reznik, D., Majka, P., Rosa, M. G. P., Witter, M. P. & Doeller, C. F. Phylogeny of neocortical-hippocampal projections provides insight in the nature of human memory. 2024.05.09.593130 Preprint at 10.1101/2024.05.09.593130 (2024).

7. Eichenbaum, H., Otto, T. & Cohen, N. J. Two functional components of the hippocampal memory system. Behavioral and Brain Sciences 17, 449–517 (1994).

8. Hebb, D. O. The Organization of Behavior; a Neuropsychological Theory. xix, 335 (Wiley, Oxford, England, 1949).

9. Duncan, K., Ketz, N., Inati, S. J. & Davachi, L. Evidence for area CA1 as a match/mismatch detector: A high-resolution fMRI study of the human hippocampus. Hippocampus 22, 389–398 (2012).

10. McClelland, J. L. & O’Reilly, R. C. Why There Are Complementary Learning Systems in the Hippocampus and Neocortex:InsightsFrom the Successesand Failuresof Connectionist Models of Learning and Memory. (1995).

11. Eichenbaum, H. & Cohen, N. J. From Conditioning to Conscious Recollection: Memory Systems of the Brain. (Oxford University Press, 2004). doi:10.1093/acprof:oso/9780195178043.001.0001.

12. Machens, C. K., Romo, R. & Brody, C. D. Functional, but not anatomical, separation of ‘what’ and ‘when’ in prefrontal cortex. J Neurosci 30, 350–360 (2010).

13. Kikumoto, A., Bhandari, A., Shibata, K. & Badre, D. A transient high-dimensional geometry affords stable conjunctive subspaces for efficient action selection. Nat Commun 15, 8513 (2024).

14. Bartolo, R., Saunders, R. C., Mitz, A. R. & Averbeck, B. B. Dimensionality, information and learning in prefrontal cortex. PLOS Computational Biology 16, e1007514 (2020).

15. Mante, V., Sussillo, D., Shenoy, K. V. & Newsome, W. T. Context-dependent computation by recurrent dynamics in prefrontal cortex. Nature 503, 78–84 (2013).

16. Harvey, C. D., Coen, P. & Tank, D. W. Choice-specific sequences in parietal cortex during a virtual-navigation decision task. Nature 484, 62–68 (2012).

17. Hedayati, S., O’Donnell, R. E. & Wyble, B. A model of working memory for latent representations. Nat Hum Behav 6, 709–719 (2022).

18. Murray, J. D., Jaramillo, J. & Wang, X.-J. Working Memory and Decision-Making in a Frontoparietal Circuit Model. J. Neurosci. 37, 12167–12186 (2017).

19. Brincat, S. L., Siegel, M., von Nicolai, C. & Miller, E. K. Gradual progression from sensory to task-related processing in cerebral cortex. Proceedings of the National Academy of Sciences 115, E7202–E7211 (2018).

20. Spens, E. & Burgess, N. A generative model of memory construction and consolidation. Nat Hum Behav https://doi.org/10.1038/s41562-023-01799-z (2024) doi:10.1038/s41562-023-01799-z.

21. Jazayeri, M. & Ostojic, S. Interpreting neural computations by examining intrinsic and embedding dimensionality of neural activity. Current Opinion in Neurobiology 70, 113–120 (2021).

22. Owen, L. L. W. & Manning, J. R. High-level cognition is supported by information-rich but compressible brain activity patterns. Proceedings of the National Academy of Sciences 121, e2400082121 (2024).

23. Linde-Domingo, J., Treder, M. S., Kerrén, C. & Wimber, M. Evidence that neural information flow is reversed between object perception and object reconstruction from memory. Nat Commun 10, 179 (2019).

24. Lifanov, J., Linde-Domingo, J. & Wimber, M. Feature-specific reaction times reveal a semanticisation of memories over time and with repeated remembering. Nat Commun 12, 3177 (2021).

25. Mirjalili, S., Powell, P., Strunk, J., James, T. & Duarte, A. Context Memory Encoding and Retrieval Temporal Dynamics are Modulated by Attention across the Adult Lifespan. eNeuro 8, ENEURO.0387-20.2020 (2021).

26. Kerrén, C., Zhao, Y. & Griffiths, B. J. A reduction in self-reported confidence accompanies the recall of memories distorted by prototypes. Commun Psychol 2, 1–11 (2024).

27. Kerrén, C., Linde-Domingo, J. & Spitzer, B. Prioritization of semantic over visuo-perceptual aspects in multi-item working memory. 2022.06.29.498168 Preprint at 10.1101/2022.06.29.498168 (2022).

28. Khodagholy, D., Gelinas, J. N. & Buzsáki, G. Learning-enhanced coupling between ripple oscillations in association cortices and hippocampus. Science 358, 369–372 (2017).

29. Jadhav, S. P., Rothschild, G., Roumis, D. K. & Frank, L. M. Coordinated Excitation and Inhibition of Prefrontal Ensembles during Awake Hippocampal Sharp-Wave Ripple Events. Neuron 90, 113–127 (2016).

30. Karimi Abadchi, J., et al. Spatiotemporal patterns of neocortical activity around hippocampal sharp-wave ripples. Elife 9, e51972 (2020).

31. Logothetis, N. K. et al. Hippocampal-cortical interaction during periods of subcortical silence. Nature 491, 547–553 (2012).

32. Joo, H. R. & Frank, L. M. The hippocampal sharp wave–ripple in memory retrieval for immediate use and consolidation. Nat Rev Neurosci 19, 744–757 (2018).

33. Norman, Y. et al. Hippocampal sharp-wave ripples linked to visual episodic recollection in humans. Science 365, eaax1030 (2019).

34. Norman, Y., Raccah, O., Liu, S., Parvizi, J. & Malach, R. Hippocampal ripples and their coordinated dialogue with the default mode network during recent and remote recollection. Neuron 109, 2767–2780.e5 (2021).

35. Vaz, A. P., Inati, S. K., Brunel, N. & Zaghloul, K. A. Coupled ripple oscillations between the medial temporal lobe and neocortex retrieve human memory. Science 363, 975–978 (2019).

36. Vaz, A. P., Wittig, J. H., Inati, S. K. & Zaghloul, K. A. Replay of cortical spiking sequences during human memory retrieval. Science 367, 1131–1134 (2020).

37. Henin, S. et al. Spatiotemporal dynamics between interictal epileptiform discharges and ripples during associative memory processing. Brain 144, 1590–1602 (2021).

38. Sakon, J. J. & Kahana, M. J. Hippocampal ripples signal contextually mediated episodic recall. Proc Natl Acad Sci U S A 119, e2201657119 (2022).

39. Dickey, C. W. et al. Widespread ripples synchronize human cortical activity during sleep, waking, and memory recall. Proc Natl Acad Sci U S A 119, e2107797119 (2022).

40. Nyhus, E. & Curran, T. Functional role of gamma and theta oscillations in episodic memory. Neurosci Biobehav Rev 34, 1023–1035 (2010).

41. Fries, P. Rhythms for Cognition: Communication through Coherence. Neuron 88, 220–235 (2015).

42. Canolty, R. T. & Knight, R. T. The functional role of cross-frequency coupling. Trends Cogn Sci 14, 506–515 (2010).

43. Sirota, A. et al. Entrainment of Neocortical Neurons and Gamma Oscillations by the Hippocampal Theta Rhythm. Neuron 60, 683–697 (2008).

44. Hyafil, A., Giraud, A.-L., Fontolan, L. & Gutkin, B. Neural Cross-Frequency Coupling: Connecting Architectures, Mechanisms, and Functions. Trends in Neurosciences 38, 725–740 (2015).

45. Fell, J. & Axmacher, N. The role of phase synchronization in memory processes. Nat Rev Neurosci 12, 105–118 (2011).

46. Colgin, L. L. Theta-gamma coupling in the entorhinal-hippocampal system. Curr Opin Neurobiol 31, 45–50 (2015).

47. Griffiths, B. J., Martín-Buro, M. C., Staresina, B. P. & Hanslmayr, S. Disentangling neocortical alpha/beta and hippocampal theta/gamma oscillations in human episodic memory formation. NeuroImage 242, 118454 (2021).

48. Canolty, R. T. et al. High Gamma Power Is Phase-Locked to Theta Oscillations in Human Neocortex. Science 313, 1626–1628 (2006).

49. Saint Amour di Chanaz, L., et al. Gamma amplitude is coupled to opposed hippocampal theta-phase states during the encoding and retrieval of episodic memories in humans. Current Biology 33, 1836–1843.e6 (2023).

50. Newman, E. L., Gillet, S. N., Climer, J. R. & Hasselmo, M. E. Cholinergic Blockade Reduces Theta-Gamma Phase Amplitude Coupling and Speed Modulation of Theta Frequency Consistent with Behavioral Effects on Encoding. J Neurosci 33, 19635–19646 (2013).

51. Daume, J. et al. Control of working memory by phase–amplitude coupling of human hippocampal neurons. Nature 629, 393–401 (2024).

52. Mormann, F. et al. Phase/amplitude reset and theta–gamma interaction in the human medial temporal lobe during a continuous word recognition memory task. Hippocampus 15, 890–900 (2005).

53. Fernández-Ruiz, A. et al. Entorhinal-CA3 Dual-Input Control of Spike Timing in the Hippocampus by Theta-Gamma Coupling. Neuron 93, 1213–1226.e5 (2017).

54. Tort, A. B. L., Komorowski, R. W., Manns, J. R., Kopell, N. J. & Eichenbaum, H. Theta-gamma coupling increases during the learning of item-context associations. Proc Natl Acad Sci U S A 106, 20942–20947 (2009).

55. Axmacher, N. et al. Cross-frequency coupling supports multi-item working memory in the human hippocampus. Proc Natl Acad Sci U S A 107, 3228–3233 (2010).

56. Axmacher, N., Elger, C. E. & Fell, J. Ripples in the medial temporal lobe are relevant for human memory consolidation. Brain 131, 1806–1817 (2008).

57. Kunz, L. et al. Ripple-locked coactivity of stimulus-specific neurons and human associative memory. Nat Neurosci 27, 587–599 (2024).

58. Staresina, B. P. & Wimber, M. A Neural Chronometry of Memory Recall. Trends Cogn Sci 23, 1071–1085 (2019).

59. Treder, M. S. MVPA-Light: A Classification and Regression Toolbox for Multi-Dimensional Data. Front. Neurosci. 14, (2020).

60. Dien, J. The ERP PCA Toolkit: An open source program for advanced statistical analysis of event-related potential data. Journal of Neuroscience Methods 187, 138–145 (2010).

61. Kobak, D. et al. Demixed principal component analysis of neural population data. Elife 5, e10989 (2016).

62. Buzsáki, G. & Draguhn, A. Neuronal oscillations in cortical networks. Science 304, 1926–1929 (2004).

63. Tamura, M., Spellman, T. J., Rosen, A. M., Gogos, J. A. & Gordon, J. A. Hippocampal-prefrontal theta-gamma coupling during performance of a spatial working memory task. Nat Commun 8, 2182 (2017).

64. Lega, B., Burke, J., Jacobs, J. & Kahana, M. J. Slow-Theta-to-Gamma Phase–Amplitude Coupling in Human Hippocampus Supports the Formation of New Episodic Memories. Cereb. Cortex 26, 268–278 (2016).

65. Schneider, M. et al. A mechanism for inter-areal coherence through communication based on connectivity and oscillatory power. Neuron 109, 4050–4067.e12 (2021).

66. Buzsáki, G. & Wang, X.-J. Mechanisms of gamma oscillations. Annu Rev Neurosci 35, 203–225 (2012).

67. Helfrich, R. F. et al. Bidirectional prefrontal-hippocampal dynamics organize information transfer during sleep in humans. Nat Commun 10, 3572 (2019).

68. Staresina, B. P., Niediek, J., Borger, V., Surges, R. & Mormann, F. How coupled slow oscillations, spindles and ripples coordinate neuronal processing and communication during human sleep. Nat Neurosci 26, 1429–1437 (2023).

69. Xiao, Z. et al. Human hippocampal ripples predict the alignment of experience to a grid-like schema. 2025.01.08.632069 Preprint at 10.1101/2025.01.08.632069 (2025).

70. Kaplan, R. et al. Hippocampal Sharp-Wave Ripples Influence Selective Activation of the Default Mode Network. Curr Biol 26, 686–691 (2016).

71. Gelinas, J. N., Khodagholy, D., Thesen, T., Devinsky, O. & Buzsáki, G. Interictal epileptiform discharges induce hippocampal-cortical coupling in temporal lobe epilepsy. Nat Med 22, 641–648 (2016).

72. Foster, D. J. & Wilson, M. A. Reverse replay of behavioural sequences in hippocampal place cells during the awake state. Nature 440, 680–683 (2006).

73. Diba, K. & Buzsáki, G. Forward and reverse hippocampal place-cell sequences during ripples. Nat Neurosci 10, 1241–1242 (2007).

74. Karlsson, M. P. & Frank, L. M. Network dynamics underlying the formation of sparse, informative representations in the hippocampus. J Neurosci 28, 14271–14281 (2008).

75. Michelmann, S. et al. Moment-by-moment tracking of naturalistic learning and its underlying hippocampo-cortical interactions. Nat Commun 12, 5394 (2021).

76. Michelmann, S. et al. Fast-timescale hippocampal processes bridge between slowly unfurling neocortical states during memory search. Preprint at 10.1101/2025.02.11.637471 (2025).

77. Reznik, D., Trampel, R., Weiskopf, N., Witter, M. P. & Doeller, C. F. Dissociating distinct cortical networks associated with subregions of the human medial temporal lobe using precision neuroimaging. Neuron 111, 2756–2772.e7 (2023).

78. Reznik, D., Margulies, D. S., Witter, M. P. & Doeller, C. F. Evidence for convergence of distributed cortical processing in band-like functional zones in human entorhinal cortex. Current Biology 34, 5457–5469.e2 (2024).

79. Heusser, A. C., Poeppel, D., Ezzyat, Y. & Davachi, L. Episodic sequence memory is supported by a theta-gamma phase code. Nat Neurosci 19, 1374–1380 (2016).

80. Vivekananda, U. et al. Theta power and theta-gamma coupling support long-term spatial memory retrieval. Hippocampus 31, 213–220 (2021).

81. Roehri, N., Bréchet, L., Seeber, M., Pascual-Leone, A. & Michel, C. M. Phase-Amplitude Coupling and Phase Synchronization Between Medial Temporal, Frontal and Posterior Brain Regions Support Episodic Autobiographical Memory Recall. Brain Topogr 35, 191–206 (2022).

82. Colgin, L. L. et al. Frequency of gamma oscillations routes flow of information in the hippocampus. Nature 462, 353–357 (2009).

83. Griffiths, B. et al. Directional coupling of slow and fast hippocampal gamma with neocortical alpha/beta oscillations in human episodic memor. Preprint at 10.1101/305698 (2018).

84. Sheng, J. et al. Higher-dimensional neural representations predict better episodic memory. Sci Adv 8, eabm3829 (2022).

85. Buzsáki, G., Leung, L. W. & Vanderwolf, C. H. Cellular bases of hippocampal EEG in the behaving rat. Brain Res 287, 139–171 (1983).

86. Parish, G., Hanslmayr, S. & Bowman, H. The Sync/deSync Model: How a Synchronized Hippocampus and a de-Synchronized Neocortex Code Memories. http://biorxiv.org/lookup/doi/10.1101/185231 (2017) doi:10.1101/185231.

87. Cayco-Gajic, N. A., Clopath, C. & Silver, R. A. Sparse synaptic connectivity is required for decorrelation and pattern separation in feedforward networks. Nat Commun 8, 1116 (2017).

88. Lanore, F., Cayco-Gajic, N. A., Gurnani, H., Coyle, D. & Silver, R. A. Cerebellar granule cell axons support high dimensional representations. Nature neuroscience 24, 1142 (2021).

89. Higgins, C. et al. Replay bursts in humans coincide with activation of the default mode and parietal alpha networks. Neuron 109, 882–893.e7 (2021).

90. Michelmann, S., Staresina, B. P., Bowman, H. & Hanslmayr, S. Speed of time-compressed forward replay flexibly changes in human episodic memory. Nat Hum Behav 3, 143–154 (2019).

91. Rigotti, M. et al. The importance of mixed selectivity in complex cognitive tasks. Nature 497, 585–590 (2013).

92. Staresina, B. P., Fell, J., Do Lam, A. T. A., Axmacher, N. & Henson, R. N. Memory signals are temporally dissociated in and across human hippocampus and perirhinal cortex. Nat Neurosci 15, 1167–1173 (2012).

93. Staresina, B. P. et al. Hippocampal pattern completion is linked to gamma power increases and alpha power decreases during recollection. eLife 5, e17397 (2016).

94. Mormann, F. et al. Latency and Selectivity of Single Neurons Indicate Hierarchical Processing in the Human Medial Temporal Lobe. J. Neurosci. 28, 8865–8872 (2008).

95. Staresina, B. P., Henson, R. N. A., Kriegeskorte, N. & Alink, A. Episodic reinstatement in the medial temporal lobe. J Neurosci 32, 18150–18156 (2012).

96. Staresina, B. P. et al. Hierarchical nesting of slow oscillations, spindles and ripples in the human hippocampus during sleep. Nat Neurosci 18, 1679–1686 (2015).

97. Stelzer, J., Chen, Y. & Turner, R. Statistical inference and multiple testing correction in classification-based multi-voxel pattern analysis (MVPA): random permutations and cluster size control. Neuroimage 65, 69–82 (2013).

98. Hörnquist, M., Hertz, J. & Wahde, M. Effective dimensionality for principal component analysis of time series expression data. Biosystems 71, 311–317 (2003).

99. Marčenko, V. A. & Pastur, L. A. DISTRIBUTION OF EIGENVALUES FOR SOME SETS OF RANDOM MATRICES. Math. USSR Sb. 1, 457 (1967).

100. Donoghue, T. et al. Parameterizing neural power spectra into periodic and aperiodic components. Nat Neurosci 23, 1655–1665 (2020).

101. Oostenveld, R., Fries, P., Maris, E. & Schoffelen, J.-M. FieldTrip: Open source software for advanced analysis of MEG, EEG, and invasive electrophysiological data. Comput Intell Neurosci 2011, 156869 (2011).

102. Bragin, A. et al. Gamma (40-100 Hz) oscillation in the hippocampus of the behaving rat. J Neurosci 15, 47–60 (1995).

103. Schreiber, T. & Schmitz, A. Improved Surrogate Data for Nonlinearity Tests. Phys Rev Lett 77, 635–638 (1996).

